# Dual-action peptide shuttles rescue Cu-amyloid-β-induced neurotoxicity and relocate Cu intracellularly

**DOI:** 10.1101/2024.09.04.611242

**Authors:** Michael Okafor, David Schmitt, Enrico Falcone, Stéphane Gasman, Laurent Raibaut, Christelle Hureau, Peter Faller, Nicolas Vitale

## Abstract

Alzheimer’s disease (AD) remains the most prevalent neurodegenerative disease characterized by intracellular neurofibrillary tangles of Tau protein and extracellular senile plaques build on Amyloid-β (Aβ) peptides. The latter result from an abnormal processing of Amyloid Precursor Protein (APP) leading to its accumulation in plaques. *Ex vivo* analyses of AD patients’ brains show an abnormally elevated concentration of metals including Cu, Zn and Fe within these plaques. Altered Cu levels have also been reported in brain regions most affected in AD. These modifications are often accompanied by reduced neuronal Cu levels and by an increased pool of extracellular labile Cu, which in turn promotes reactive oxygen species (ROS) formation. To counteract this Cu dyshomeostasis and limit Cu-Aβ-induced extracellular ROS generation, we designed and synthesized two Cu(II)-selective peptide shuttles based on kinetically optimized ATCUN sequences for fast Cu(II) extraction out of Aβ: DapHH-αR5W4^NBD^ and HDapH-αR5W4^NBD^. They were also equipped with a fluorophore that showed a very strong response to Cu(II)-binding and release. Interestingly, these two Cu(II) shuttles displayed a dual mode of action. They promptly retrieve Cu from extracellular Aβ, stop the associated ROS formation, and hence protect both cell culture models and organotypic hippocampal slices (OHSCs) from Cu(Aβ)-induced neurotoxicity. Moreover, these shuttles import and redistribute bioavailable Cu inside cells with a sequence-dependent kinetics.

## Introduction

Copper (Cu) is an essential that must be tightly regulated and correctly distributed within living organism to sustain a large number of biological mechanisms^1–3^. Disturbance in its distribution can cause severe disorders including Menkes and Wilson’s disease as well as neurodegenerative disorders such as Alzheimer’s disease (AD)^1,4,5^. The loss of Cu homeostasis can lead to dysregulation of normal cuproprotein functions such as antioxidant defense, iron (Fe) homeostasis or mitochondrial energy conversion^2^. In AD patients, there is an increase in blood non-ceruloplasmin bound Cu^6–10^. Interestingly, a 3-4 fold higher risk of AD has been associated with the presence of excess Cu in the blood^10^. Additionally, patients with the highest concentrations of Cu not bound to ceruloplasmin perform worse on neuropsychological tests^10^. Equally, results from several groups have shown a general decrease in brain Cu levels in AD using various quantitative approaches^6,7^. Similarly, recent studies have shown a decrease in Cu levels in the cytoplasm of brain cells, which is associated with a 25% increase of exchangeable Cu^11,12^.

An aberrant decrease of Cu in a biological compartment often leads to loss of function for cuproproteins. Equally, an increase in Cu concentration in a compartment most likely leads to the binding of Cu to inappropriate sites on proteins/ligands, potentially inducing a toxic gain of function. As such, the increase in extracellular labile Cu favors its binding to Aβ peptides, driving toxicity in AD^11–14^. The resulting Cu(Aβ) complex catalyzes the aerobic formation of reactive oxygen species (ROS) in the presence of a physiological reductant (e.g. ascorbate, AscH^-^) by cycling between Cu(I) and Cu(II)^15,16,17^. This reaction is possible because the Aβ peptide forms a complex with both Cu(II) (dissociation constant, K_d_ ≈ 10^-9^-10^-10^ M) and Cu(I) (K_d_ ≈ 10^-7^-10^-10^ M) at neutral pH^18^, following a highly intricate redox mechanism^19^. Cu(Aβ) complexes have therefore been proposed to contribute to the oxidative stress described in AD^20^. This stress takes various forms, ranging from peroxidation of cell membranes, oxidation of DNA and mitochondria^17,21^. ROS can also generate oxidatively-modified Aβ_1-40/42_ peptides, which is linked to a modified aggregation propensity or an increased resistance to proteolytic degradation ^22,23^.

Cu disbalance thus seems to play an important role in AD progression and restoring its homeostasis might be important for disease remediation. Therefore, in complement to chelating this excess extracellular Cu, it is of interest to redistribute Cu inside neuronal cells where it is missing, in a “one stone, two birds” approach. This drove the design of ionophores that act as transmembrane transporters to shuttle metal ions, such as Clioquinol (iodochlorhydroxyquin, CQ) and its derivative, PBT2, as well as GTSM (glyoxal-bis(4-methylthiosemicarbazonato))^24–27^. These molecules chelate Cu(II) and Zn(II) to form complexes capable of crossing cell membranes. They have shown convincing reduction in metal-modulated Aβ aggregation *in vitro*, accompanied by improved performances in various cognitive tests on AD murine models^27,28^. CQ reached Phase II in clinical trial, showing improvement in various cognitive tests in advanced AD patients^29^, but negative side-effects with prolonged treatment halted further development^30,31^. PBT2 developed later as a safer CQ analog was proved ineffective in phase II clinical trial^32^. Therefore, Cu(II)-selective ionophores with appropriate Cu(II) affinity and selectivity in the context of AD remain to be developed. Importantly, many available Cu(II) ionophores such as GTSM, Clioquinol or PBT2 exhibit appropriate Cu(II) affinities but suffer from poor selectivity in biological conditions^24,33,34^.

The recently developed AKH-αR5W4^NBD^ shuttle was shown to import bioavailable Cu into cells^35^. Its Cu(II)-selective binding domain of this shuttle is an Amino Terminal Cu(II)-and Ni(II)-Binding (ATCUN) motif, a tripeptide motif with His residue in the third position (H_2_N-XxxZzzHis). ATCUN motifs are known to be very selective for Cu(II) versus Zn(II), the latter being far more abundant than Cu(II) in the brain, thus making this scaffold very appropriate for selective Cu(II) targeting^36,37^. αR5W4, a Cell Penetrating Peptide (CPP) optimized for efficient membrane penetration^38^, was chosen as an agent to allow cell delivery of the Cu. Additional studies indicated that AKH-αR5W4^RhodB^ internalization in HeLa and PC12 cells proceeds via ATP-dependent endocytosis pathways, such as clathrin-mediated endocytosis, caveolin-mediated endocytosis or micropinocytosis, therefore preventing intracellular oversaturation^39^. However, this shuttle lacked the kinetics to quickly retrieve Cu from Aβ, thus hampering its effectiveness and neuroprotective potential. In this work, we optimized the ATCUN motif of AKH-αR5W4^NBD^ was kinetically optimized for the ability to accelerate Cu(II) retrieval from Aβ and suppress Cu(Aβ) induced ROS production.

## Materials and methods

### Materials

All compounds used during this study were purchased from accredited merchants including Sodium L-ascorbate (Sigma, A4034), DMEM/F-12 (Gibco, 11320033), Dulbecco′s Modified Eagle′s Medium - high glucose (Sigma, D5796), MEM (Thermofisher, 12360038), Horse serum (Gibco, 26050070), Fetal calf serum (Gibco, 10270-160), Penicillin/streptomycin (Sigma, P4458), Trypsin (Gibco, 25300-054), MTT (Fisher Scientific, 10133722), Anti-Giantin antibodies (Abcam, ab24586), Monoclonal Anti-ATP7A antibodies (Abcam, ab13995), Rabbit anti-Mouse Alexa Fluor 555 (Invitrogen, A-21427), Donkey anti-Chicken Alexa Fluor 488 (Invitrogen, A78948), Recombinant Mouse Anti-Iba1 (Abcam, ab283319), Monoclonal Rabbit Anti-CD68 (Abcam, ab125212) and Hoechst (Thermofisher, 62249).

### Shuttle Synthesis and Purification

Peptide synthesis was carried out according to a general SPPS protocol^35^. In short, peptide synthesis was performed using Biotage® Initiator+ Alstra™. Fmoc-Rink-Amide Tenta XV RAM resin (loading 0.24 mmol.g^-1^) was used. 0.5 M 1: 1 ethyl cyano(hydroxyimino)acetate (oxime)/diisopropylcarbodiimide (DIC) was used for activation, and 2 M DIEA dissolved in N-methyl-2-pyrrolidone (NMP) was used as base. 5 M acetic anhydride was used to cap unreacted free amino groups. The reaction time was 2 × 30 minutes coupling at ambient temperature and 2 × 5 min coupling using microwaves (μWF) at 75 °C with the exception of basic amino acids which were coupled for 2 x 30 min. Fmoc deprotection was carried out using μWF at 75 °C. Dap (2,3-diaminopropionic acid) was added as the Nε-allyloxycarbonyl-protected (Fmoc-Dap (Alloc)–OH) form to allow specific sidechain deprotection in order to graft the fluorophore NBD. The Alloc protecting group of the Dap side chain is deprotected using tetrakis(triphenylphosphine)palladium(0), which is a palladium-based mechanism^40^. The NBD fluorophore is added to the free lysine by nucleophilic substitution of 4 eq. NBD-Cl in the presence of DIEA. Finally, deprotection was carried out using TFA in the presence of scavengers (H_2_O and triisopropylsilane). Purification was performed using a HPLC with reverse-phase analytical (Hitachi Primaide equipped with a column XBridge C18 BEH 300 Å, 5 μm, 4.6 × 150 mm, 37 °C), the purity and mass were analysed by LC-MS (LTQ-XL Thermo).

### Aβ monomer preparation

Aβ_1-40/42_ peptides (DAEFRHDSGYEVHHQKLVFFAEDVGSNKGAIIGLMVGGVV-IA) were purchased from GeneCust (Dudelange, Luxembourg) with a purity grade > 95%. Three different batches were obtained for Aβ_1-40_ and 1 batch for Aβ_1-42_. They were solubilized in 100% HFIP (Hexafluoro-2-propanol) for 1 h under sonication to give a concentration of ≈40 g/L (≈1 mM). Afterwards, 100 µL was placed in 0.5 mL Eppendorf tubes and left under a fume hood for 1 week to completely evaporate HFIP. After evaporation, the Aβ peptides formed a film that was conserved at-20°C until usage. Before use, Aβ film was solubilized in 20 µL DMSO, vortexed briefly, then centrifuged for 30 seconds and then re-vortexed. UV-Vis measurement was carried by adding 0.5 µL of Aβ stock solution in 100 µL Milli-Q water (ρ =18.2 MΩ⋅cm-1). The absorbance at 280 nm was obtained and using the molar extinction coefficient of 1 400 M^-1^ cm^-1^ for Aβ monomers^41^, the final theoretical concentration of Aβ was ≈ 5 mM.

### Peptide shuttle titration via Cu

In 96-well plates, peptide shuttles were added to wells containing 100 mM HEPES to a final concentration of 4 µM. Then an eq of Cu(II) was added by step of 0.1 eq. The fluorescence was followed using CLARIOstar (BMG LABTECH) plate reader at an excitation wavelength of 477 nm and the emission at 545 nm in a UV-STAR® MICROPLATE, 96 well half area microplate.

### Selective Cu(II) retrieval from media by peptide shuttle

In 96-well plates, 4 µM Cu(II) was added to wells containing either solutions of 100 mM HEPES or DMEM:F12 Nutrient 1:1, and an eq concentration of peptide or Cu(II) or Cu(II) + Mn(II), Zn(II) and Fe(III). The fluorescence was followed using CLARIOstar (BMG LABTECH) plate reader at an excitation wavelength of 477 nm and the emission between 490 and 600 nm for 2h at an interval of 5min in a UV-STAR® MICROPLATE, 96 well half area microplate.

### Retrieval of Cu by peptide shuttles from Aβ

The emission intensity of NBD on peptide shuttles was used as readout. Using a Horiba Fluoromax plus fluorimeter, in a 500 µL 1 cm path quartz cuvette, 10 µM Aβ and 5 µM AKH-αR5W4^NBD^, DapHH-αR5W4^NBD^ or HDapH-αR5W4^NBD^ in a solution of 100 mM HEPES or 10% DMEM was recorded at 545 nm to determine the estimated 100% unsaturated emission intensity. The same was done for 10 µM Cu(II)-Aβ 1:2 and 5 µM Cu(II)-AKH-αR5W4^NBD^, DapHH-αR5W4^NBD^ or HDapH-αR5W4^NBD^ 1:1, to give the estimated 100% saturated emission level. Afterwards, 5µM of AKH-αR5W4^NBD^, DapHH-αR5W4^NBD^, or HDapH-αR5W4^NBD^ was added to the mixture of 10µM Cu(II):Aβ 1:2, the emission at 545 nm was recorded every 30 seconds for up to 2h at 25°C.

### Release of Cu by peptide shuttles in GSH

Using a Horiba Fluoromax plus fluorimeter, in a 500 µL 1 cm path quartz cuvette, 5 mM GSH and 5µM AKH-αR5W4^NBD^, DapHH-αR5W4^NBD^, or HDapH-αR5W4^NBD^ was added to a 100 mM HEPES buffer pH 7.4. The solution was excited at 477 nm and recorded at 545 nm, this gives the emission intensity of the peptide without Cu(II). The intensity of Cu(II)-AKH-αR5W4^NBD^, DapHH-αR5W4^NBD^ or HDapH-αR5W4^NBD^ was recorded in the absence of GSH, then 5 mM GSH was added from a stock solution at 200 mM and left to react for up 12h at 37°C.

### Inhibition of Ascorbate oxidation by peptide shuttles

Using CLARIOstar (BMG LABTECH) plate reader, experiments were carried out in a UV-STAR® MICROPLATE, 96 WELL, HALF AREA microplate. The absorbance of 100 µM AscH^-^ (OD ≈ 1.5) at 265 nm was followed under different conditions to indirectly follow ROS production. The final volume per well was 100 µL and was buffered at pH 7.4 using 100 mM HEPES. The initial rate of ascorbate consumption was determined by taking the first 20 min portion of the curve considered as linear.

### Toxicity assay on PC12 cells

PC12 cells were maintained in culture in a DMEM high glucose (4500 mg/mL) with 10% horse serum, 5% FBS and 1% penicillin/streptomycin as described previously^42^. The cells were split once a week using trypsin for detachment and replaced by a fresh batch of cells after a maximum of 15 splits.

In 96 well plates, 50 000 cells were plated and incubated for 24h. A solution of monomerized Aβ complex in 10% DMEM (diluted in a salt solution (0.2g/L CaCl_2_, 0.0001g/L Fe(NO_3_)_3_, 0.098g/L MgSO_4_, 0.4g/L KCl, 3.7g/L NaHCO_3_, 6.4g/L NaCl, 0.11g/L NaH_2_PO_4_, 4.5g/L D-Glucose)) was prepared in microcentrifuge tubes, followed with/without addition of AKH-αR5W4^NBD^, DapHH-αR5W4^NBD^ or HDapH-αR5W4^NBD^ for 10 min or 1h. Afterwards, 100 µM AscH^-^ was added to the tubes and vortexed, the mixture was then added to PC12 cells. After 24h, the cell media was replaced with 200 µL of pure DMEM containing 10% of 5 mg/mL MTT solution and incubated for 4 h at 37°C. After a 4 h incubation at 37°C the medium was replaced with 150 µL DMSO and agitated for 5 min at room temperature. In a new 96 well plates, 50 µL solution of each well was diluted in 150 µL of pure DMSO solution. Lastly, the absorbance was read at 595 nm. Each condition corresponds to the average of three recordings on the same plate and each experiment was replicated independently three times.

### ATP7A delocalization assay

Glass coverslips were treated with Poly-L-Lysine for 15 min in 4-well NUNC plates. Afterwards, the coverslips were rinsed with enriched DMEM media. PC12 cells were then cultured at a density of 1 x 10^5^ per well, 24 h before experiments. In 10% DMEM solution, was added EDTA, GTSM or Cu shuttles alone or precomplexed to Cu(II). This mixture was added to the cells and incubated for 1 h. After the experiment, the medium was removed, and cells were fixed with 4% PFA (paraformaldehyde) then permeabilized and saturated in 0.2% Triton X-100 and 1X PBS containing 10% DS (Donkey serum) at 37°C. Cells were then incubated overnight at 4°C with chicken anti-mouse ATP7A 1/5000 and rabbit anti-mouse Giantin 1/500 in 3% BSA (Bovine Serum Albumin). The next day the cells were washed 3 times with PBS and then incubated with donkey anti-chicken AlexaFluor 488, anti-rabbit AlexaFluor 555 secondary antibodies and Hoechst at 1/1000 dilution for 45 min at 37°C before being rinsed and mounted in Mowiol.

### Organotypic hippocampal slice culture (OHSC)

The procedure recapitulates that of Stoppini et al. 1991 with minor modifications^43^. CD1 mice between ages P6 and P10 were used for OHSC. The dissection media consists of cold Hanks balanced salt solution (HBSS) and the llice culture media consists of 50% MEM solution, 25% HBSS, 25% Horse serum inactivated, 1% Strep-penicillin, 25 mM HEPES and 2 mM L-Glutamine. D-Glucose was added to the cell culture solution for a final concentration of 4.5 g/L (the Culture media should be at a pH 7.2).

Mice were decapitated and placed immediately in a 25 mL beaker containing ice cold HBSS. Using scissors, the skull was cut through and the brain extracted, then transferred to a 30 mm petri dish filled with cold HBSS. The hippocampus was extracted by holding down one hemisphere with forceps and separating the other with a spatula. Both hippocampi were extracted and placed on the Mcllwain tissue chopper board perpendicular to the blade, with the micrometer set at 400 µm and the blade wetted with cold HBSS before chopping. After brain slices were ready, the slices were transferred to a 30 mm petri dish filled with cold HBSS, and each slice was separated using 2 spatulas. Each slice containing the dentate gyrus, CA1, and CA3 regions, were placed on the Millicell insets with 6 hippocampal slices per well of a 6-well plate. The plate was placed in an incubator and the media was changed every 2-3 days. The slices were kept in culture for two weeks, to acclimatize to their new environment before experiment.

### Cu toxicity rescue experiment OHSCs

After two weeks in culture, hippocampal slices were treated with different conditions for 48h in DMEM solution containing 25 mM HEPES. Prior to the experiment, preparation of Aβ monomers for monomeric conditions was carried out (see supplementary materials and methods).

Cu(II)Aβ complex was made by mixing 0.5 eq Cu (10 µM) with 1 eq Aβ (20 µM). After complexation of Cu(Aβ), DapHH-αR5W4^NBD^ or HDapH-αR5W4^NBD^ was added or not at 0.5 eq and vortexed. The solution was immediately placed on top of the slices (with time solution will drain into the well out of the Millicell inset). Slices were left for 48h at 37°C.

### PI Staining

A salt solution (0.2g/L CaCl_2_, 0.0001g/L Fe(NO_3_)_3_, 0.098g/L MgSO_4_, 0.4g/L KCl, 3.7g/L NaHCO_3_, 6.4g/L NaCl, 0.11g/L NaH_2_PO_4_, 4.5g/L D-Glucose) containing 5 µM PI and 25 mM HEPES was prepared, and 100 µL was placed in each well on top of every slice and incubated for 1h at 37°C. Afterwards, slices were fixed with 4% PFA for 40 min. From each condition, 3 slices were cut out of the millicell insert after fixation and treated with Hoechst for 30 min and then mounted on slides. Slides were left at room temperature to semi-dry for 1 h and then images were acquired at 405 nm and 555 nm using a fluorescence microscope, ZEISS Axio Imager M2 coupled with a motorized board for mosaic imaging.

### Immunohistochemistry

The remaining 3 slices were cut out of the millicell insert and were washed in PBS. The slices were permeabilized and nonspecific sites were blocked in PBS solution containing 0.3% triton and 10%

Goat serum for 3 hours under agitation. Afterwards, slices were rinsed once in PBS and then incubated in PBS containing 1 mg/ml BSA (bovine serum albumin) overnight at 4°C with primary antibodies (Mouse Anti-Mouse Iba1 1:100 and Rabbit Anti-Mouse CD68 1:1000). Slices were washed 3 times in PBS for 5 min each with gentle shaking, and then incubated with secondary anti-mouse 555 nm and anti-rabbit 647 nm antibodies at 1:1000 dilution in PBS containing 1 mg/mL BSA for 3 hours at room temperature with gentle shaking. Finally, slices were washed 3 times in PBS for 5 min each with gentle shaking and mounted on glass slides by placing the slices with the membrane side flat on the slides. Mowiol was added on each slice followed by the cover. Slides were left to dry at room temperature in obscurity for at least 24h before confocal imaging using Leica SP5.

### Analysis of Cell viability OHSC

Viability measurements were made by cell counting total cell number via the nucleus staining by Hoechst using the QuPath software. Cells were also incubated with propidium iodine (PI) and the mean fluorescence intensity of the PI staining was calculated in each cell nucleus. Cells with PI staining above a threshold were deemed PI positive and divided by the total cells (Hoechst positive) to give the percentage of Hoechst+ PI+ cells.

### Analysis of Microglia activation OHSC

Microglia activation analysis was carried out by labelling cells positive with both Iba1 and CD68. Iba1 cells were counted using QuPath software and labelled as microglia cells. From cells positive for Iba1, the average intensity of the microglial cells with a threshold of CD68 labelling were labelled as activated microglial cells. The percentage of activated microglial cells are expressed as CD68 positive cells divided by Iba1 positive cells. All data were normalized in respect to the control group.

## Statistical analysis

All analyses were carried out using GraphPad prism 9.3.1. Data set underwent Shapiro test for normality. For comparisons of the mean, parametric test with Tukey Multiple Comparison post-test or non-parametric Kruskal–Wallis with Dunn’s multiple comparisons test.

## Results

In the quest for efficient peptide shuttles, in regards to the arrest of ROS production, the rate of Cu(II) removal from Cu(Aβ) complexes was identified as an additional key parameter^35^. Based on a large series of kinetic investigations, the presence of two His, either at positions 1 and 3 (H_2_N-HisXxxHis) or positions 2 and 3 (H_2_N-XxxHisHis), was shown to allow the ATCUN scaffold to overcome such kinetic limitation^44,45^. Therefore, we designed, synthesized and probed two different ATCUN motifs. These two shuttles H_2_N-Dap-His-His-αR5W4^NBD^ (DapHH-αR5W4^NBD^) and H_2_N-His-Dap-His-αR5W4^NBD^ (HDapH-αR5W4^NBD^), were functionalized with a nitrobenzodiazole (NBD) fluorophore, on the Dap (2,3-diaminopropionic acid, a lysine analogue with a shorter side chain) residue inside the ATCUN motif, for closer proximity to the bound Cu(II) (see Methods and Figure S1A-C).

### Physico-chemical characterizations of Cu(II) peptide shuttles

The differences in the physico-chemical properties of the Cu(II)-selective peptides shuttles (DapHH-αR5W4^NBD^ and HDapH-αR5W4^NBD^) in comparison to the parent molecule, AKH-αR5W4^NBD^, were investigated (Figure 1A, S1A-E). The UV-Vis spectra display similar absorption bands at 350 and 490 nm due to the NBD chromophore (Figure S1D). A red-shift was observed for the Cu(II)-HDapH-αR5W4^NBD^ complex (compare Figure S1D and S1E, red traces), suggesting a structural change upon Cu(II)-binding. The emission of both DapHH-αR5W4^NBD^ and HDapH-αR5W4^NBD^ was then studied by fluorimetry. Since the NBD fluorophore has an excitation maximum at 480 nm and an emission maximum at 545 nm in aqueous medium, the two peptides were excited at 477 nm and the fluorescence emission spectra recorded between 490 nm and 600 nm. The emission maximum was observed at 545 nm and the successive additions of Cu(II) in a 0.1 equivalent step led to a linear decreased in emission intensity at 545 nm up to 1 equivalent (Figure S2A). For both DapHH-αR5W4^NBD^ and HDapH-αR5W4^NBD^, 98% quenching of NBD emission was recorded thus validating the intended design of the peptides to display a high percentage of fluorescence quenching upon Cu(II)-binding.

**Figure 1:**
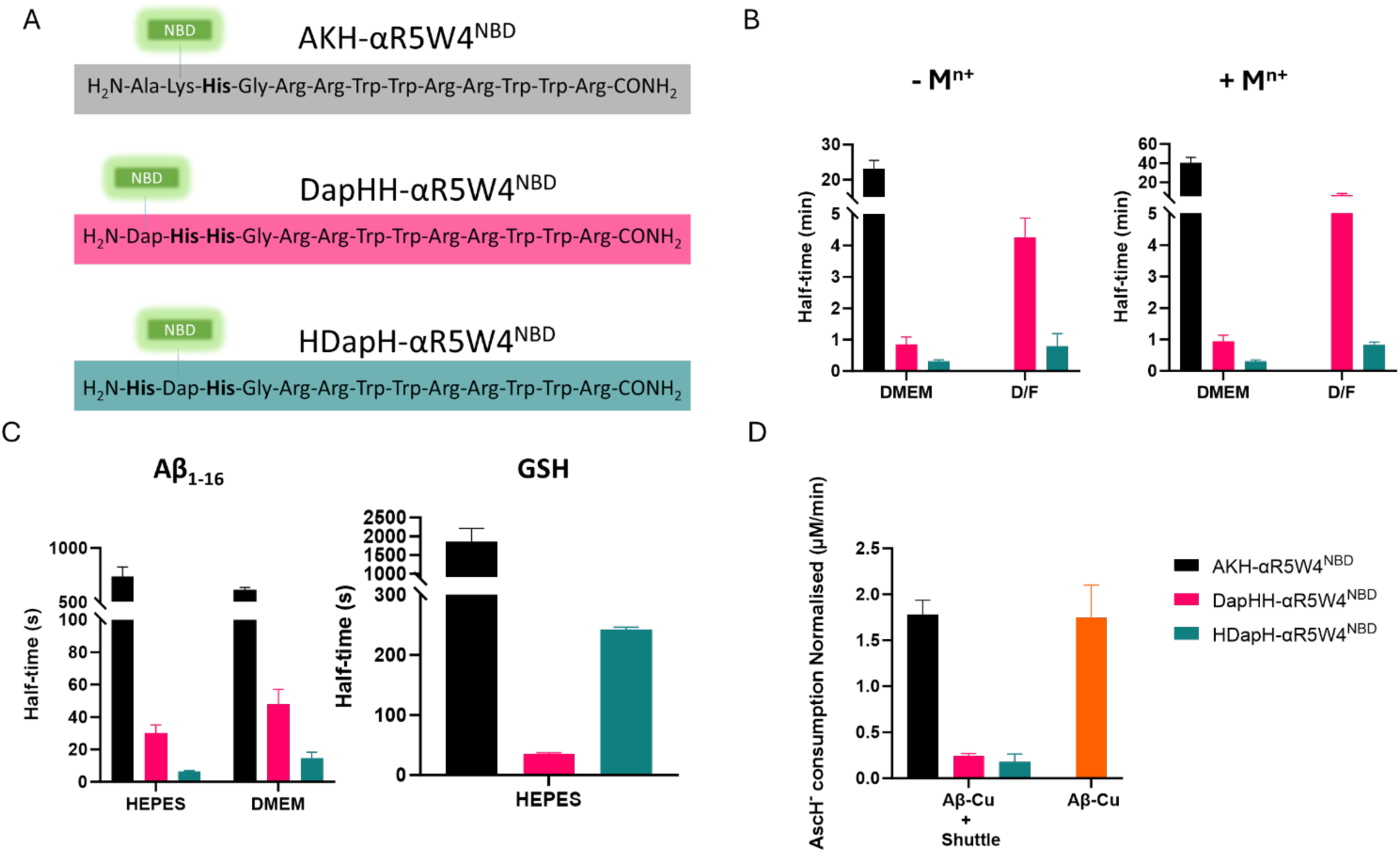
Properties of DapHH-αR5W4^NBD^ and HDapH-αR5W4^NBD^ shuttles. (**A**) Amino acid sequence of the peptide shuttles under study: AKH-αR5W4^NBD^, DapHH-αR5W4^NBD^ and HDapH-αR5W4^NBD^. (**B**) Half-time of the selective withdrawal of Cu(II) by the shuttles in cell culture media containing Cu(II) alone or in addition to M^n+^ (Fe (III), Zn(II) and Mn(II)). AKH-αR5W4^NBD^ Cu(II) withdrawal was too slow in D/F media, data not shown. Conditions: Cu(II)=AKH-αR5W4^NBD^=DapHH-αR5W4^NBD^=HDapH-αR5W4^NBD^= 5 µM, DMEM 10%, D/F 100%, 25°C, n=3. **(C)** Retrieval of Cu(II) from Aβ_1-16_ by the shuttles in HEPES or DMEM 10% media and reduction/release of Cu(II) by GSH in HEPES. Conditions: AKH-αR5W4^NBD^ >DapHH-αR5W4^NBD^=HDapH-αR5W4^NBD^= 5 µM; Aβ_1-16_ =10 µM; Cu(II)=5 µM, HEPES 100 mM pH 7.4, 25°C, n=3. **(D)** Inhibition of AscH^-^ consumption by the shuttles monitored by the absorbance of AscH^-^ at 265 nm (see Figure S2E for the complete kinetic curves). Addition of AscH^-^ was carried out at t_0_ and Cu(II)Aβ_1-16_ at 5 min and addition of the shuttles at 10 min. Graph shows the initial consumption of AscH^-^ in the first 20 min after addition of the shuttles in the presence of Cu(II)Aβ_1-16_, normalized with values from samples in the absence of Cu(II)Aβ_1-16_. Conditions: AscH^-^= 100 µM, 5 µM Cu(II)DapHH-αR5W4^NBD^ or Cu(II)HDapH-αR5W4^NBD^, Aβ_1-16_= 10 µM, Cu(II)= DapHH-αR5W4^NBD^= HDapH-αR5W4^NBD^ = 5 µM, HEPES 100 mM pH 7.4, n=2 independent experiments.

Next, we investigated the capacity of the shuttles to withdraw Cu(II) from cell culture media, DMEM 10% (DMEM diluted in salt solution) and DMEM/F12 (D/F). For this, we spiked DMEM 10% media or D/F media with 5 µM Cu(II) for one minute, then added AKH-αR5W4^NBD^, DapHH-αR5W4^NBD^ or HDapH-αR5W4^NBD^ and then followed the NBD emission for 5h in a plate reader. Controls were carried out simultaneously in the absence of Cu and with the shuttles pre-complexed with Cu(II) before addition to the media, allowing the estimation of NBD emission at both the 0% saturated states and the 100% saturated states, respectively (Figure 1B, S2B-E). Both DapHH-αR5W4^NBD^ and HDapH-αR5W4^NBD^ out-performed AKH-αR5W4^NBD^ in the rate of withdrawing Cu(II) from the cell culture media (Figure 1B). Indeed, all three peptide shuttles were capable of withdrawing Cu(II) from DMEM 10% media, with the rate following the trend; HDapH-αR5W4^NBD^ > DapHH-αR5W4^NBD^ > AKH-αR5W4^NBD^ (Figure S2B&D). In addition, HDapH-αR5W4^NBD^ completely withdrew Cu(II) from D/F media (higher concentration of amino acids), with DapHH-αR5W4^NBD^ retrieving >80% and AKH-αR5W4^NBD^ withdrawing ≈50% over the measurement duration (Figures 1B, S2C&E and Table S1). Altogether, these results confirm the successful design of shuttles with faster Cu(II)-retrieval kinetics.

We then validated the selectivity of DapHH-αR5W4^NBD^ and HDapH-αR5W4^NBD^ to withdraw specifically Cu(II) from cell media containing other essential d-block metal ions. For this, we spiked both DMEM 10% and D/F media with Cu(II) and M^n+^ (Fe(III), Zn(II) and Mn(II)), each at 5 µM before addition of AKH-αR5W4^NBD^, DapHH-αR5W4^NBD^ or HDapH-αR5W4^NBD^ after a minute. No major difference in the percentage or withdrawal kinetics for the three peptide shuttles was observed in the presence or absence of other d-block metal ions in DMEM was observed (Figures 1B, S3A-D and Table S1). However, a slight reduction in the average kinetics of Cu withdrawal by DapHH-αR5W4^NBD^ was observed in D/F, although HDapH-αR5W4^NBD^ stayed constant. Again, HDapH-αR5W4^NBD^ was generally the fastest followed by DapHH-αR5W4^NBD^ and then AKH-αR5W4^NBD^ (Figure 1B).

We next compared the kinetics of the selective Cu(II) withdrawal by DapHH-αR5W4^NBD^ and HDapH-αR5W4^NBD^ with that of GTSM, a prototypical ionophore (Figure S4). HDapH-αR5W4^NBD^ but not DapHH-αR5W4^NBD^ displayed comparable kinetics of Cu(II) withdrawal in comparison to that of GTSM in both DMEM 10% and D/F media (half-time <1 min).

Next, we investigated the capacity of the two peptide shuttles to withdraw Cu(II) from Aβ_1-16_, the domain responsible for Cu binding in Aβ peptides^22,46^. Rapid retrieval of Cu(II) from Aβ_1-16_ was observed in HEPES buffer in 10% DMEM for both peptide (Figure 1C, S5A,B), with the complete retrieval occurring in less than two minutes compared to over an hour for AKH-αR5W4^NBD^. Among these two shuttles, HDapH-αR5W4^NBD^ acted faster than DapHH-αR5W4^NBD^ (Figure 1C).

We then investigated whether physiological levels of glutathione (GSH, 5 mM) could reduce Cu(II) bound to DapHH-αR5W4^NBD^ and to HDapH-αR5W4^NBD^, given that reduction of Cu(II) by a reducing agent like GSH renders bound Cu bioavailable. Both DapHH-αR5W4^NBD^ and HDapH-αR5W4^NBD^ displayed half-times of only a few minutes, whereas that of AKH-αR5W4^NBD^ required tens of minutes (Figure 1C, Figure S5C and Table S2). Notably, Cu(II)-DapHH-αR5W4^NBD^ was reduced faster than Cu(II)-HDapH-αR5W4^NBD^ (Figure 1C, Figure S5C and Table S2).

We next tested the capacity of the shuttles to inhibit Cu(Aβ)-induced ROS production catalyzed by AscH^-^. AscH^-^ was monitored at 265 nm, a classical assay that mirrors ROS formation^35^. The initial rate of ROS production by Cu(II)Aβ_1-16_ indirectly measured during the first 20 minutes after addition of AscH^-^ was 1.6 ± 0.1 µM/min, in agreement with reported values^18^. Upon addition of DapHH-αR5W4^NBD^ or HDapH-αR5W4^NBD^, an immediate reduction in ROS production was observed (Figure 1D and S2D,E), which was not the case for AKH-αR5W4^NBD^ shuttle^35^.

In summary, both DapHH-αR5W4^NBD^ and HDapH-αR5W4^NBD^ have proven to be much faster than AKH-αR5W4^NBD^ in retrieving Cu(II) from Aβ_1-16_ and therefore in suppressing Cu(Aβ)-induced ROS production. In the following studies, we investigated whether these shuttles are also able to transfer and release Cu intra-cellularly in addition to preventing cells from extracellular Cu(Aβ)-induced ROS toxicity.

### Cellular characterization of DapHH-αR5W4^NBD^ and HDapH-αR5W4^NBD^ in PC12 cells

Firstly, we tested their toxicity of these shuttles on PC12 cells, a well-known neurosecretory cell model^47^. Neither DapHH-αR5W4^NBD^, HDapH-αR5W4^NBD^ nor the corresponding Cu(II) complexes were toxic bellow 10 µM (Figure S6A). This indicates that micromolar concentrations of the peptide shuttles, unlike GTSM (toxic at nanomolar concentration), do not lead to intracellular toxic levels of Cu (Figure S6B). It is possible that the cell toxicity induced by GTSM results from too high levels, too fast release, and/or release of Cu in inappropriate intracellular compartments. We then investigated the cell entry of 5 µM of these Cu(II)(peptides) complexes into PC12 cells using confocal microscopy. Over a 30 min incubation, we observed membrane binding followed by a gradual increase in intracellular fluorescence attributed to an intracellular release of Cu by the peptide shuttles (Figure 2A and S7A). Compared to the peptide shuttle without Cu, the Cu(II)-complexed shuttles showed lower membrane and intracellular fluorescence within the first minutes of incubation, likely reflecting a delay for the Cu(II) expulsion from the Cu(II)(shuttle) complexes after internalization (Figure 2A). The intracellular fluorescence signal mostly corresponded to vesicular structures similar to endosomes, which points to a cell entry mechanism involving endocytosis as recently established for the αR5W4 CPP^37^. Interestingly, cells incubated with Cu(II)(HDapH-αR5W4^NBD^ displayed a more rapid increase in vesicular fluorescence for compared to those incubated with Cu(II)(DapHH-αR5W4^NBD^). This was unexpected given that *in vitro* data demonstrated that Cu(II)(DapHH-αR5W4^NBD^) is reduced faster than Cu(II)(HDapH-αR5W4^NBD^) (Figure 1C, GSH). This result suggests that Cu release from the shuttles occurs through mechanisms other than the reductive environment of intracellular compartments.

**Figure 2:**
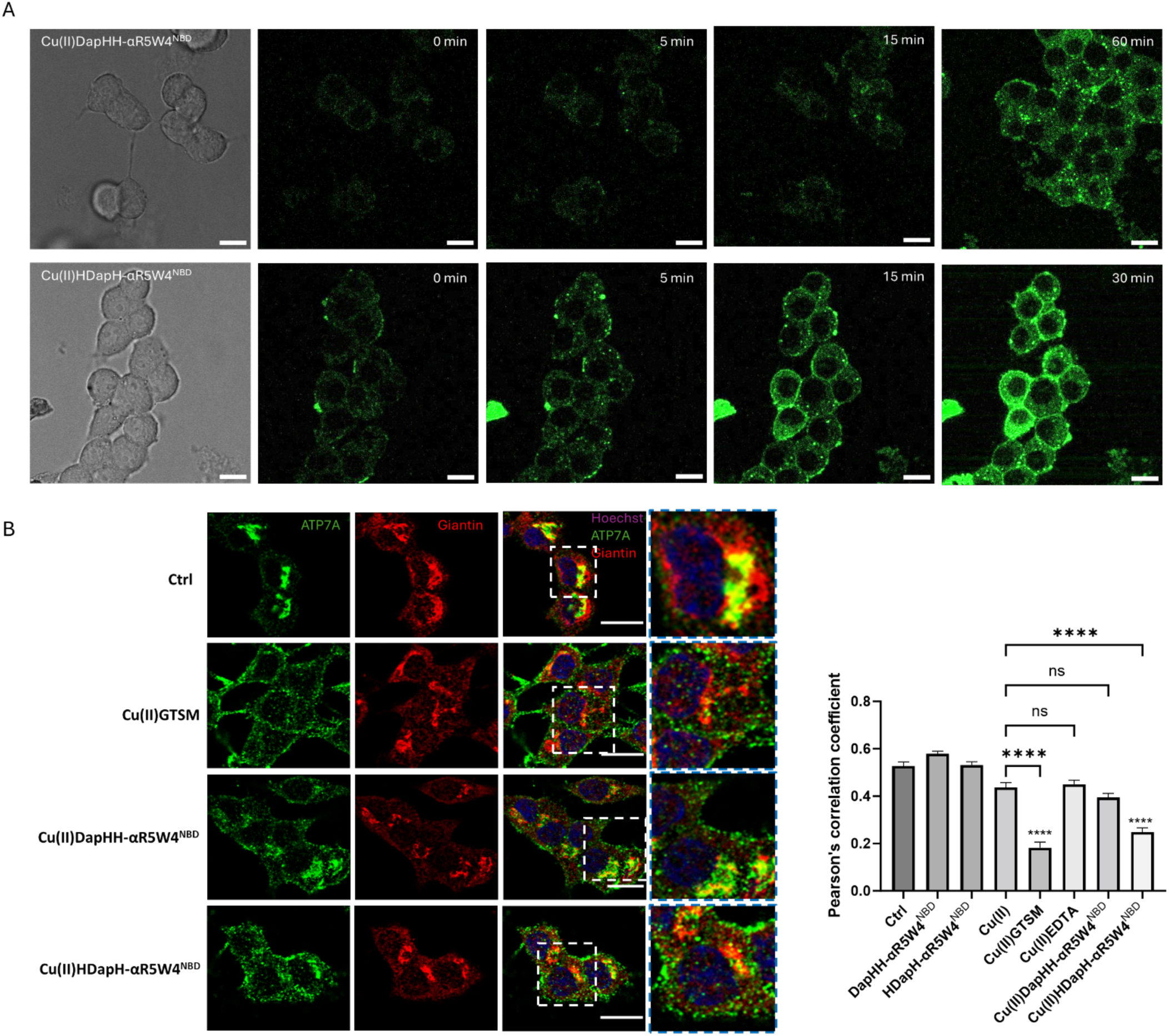
Cu import by HDapH-αR5W4^NBD^ induced delocalization of ATP7A from the TGN to vesicular structures.(A) Representative images at the indicated time obtained from live imaging performed on 5 x 10 ^4^ PC12 cells incubated with 5 µM Cu(II)(DapHH-αR5W4^NBD^) or Cu(II)(HDapH-αR5W4^NBD^), at 37°C using a Leica TCS SP5 (II) confocal microscope, with an excitation wavelength at 477 nm for 30 min. Time point zero is about 1 minute after incubation with the peptide shuttles. Bars = 10 µm. **(B)** Quantification of the colocalization between ATP7A and Giantin (Golgi marker) staining was measured as Pearson’s correlation coefficient. PC12 cells were incubated for 1h in DMEM media alone (control = Ctrl) or, 5 µM Cu(II), Cu(II)-EDTA, DapHH-αR5W4^NBD^, Cu(II)(DapHH-αR5W4^NBD^), HDapH-αR5W4^NBD^ and Cu(II)(HDapH-αR5W4^NBD^), or 1 µM Cu(II)-GTSM, before fixation and process for immunostaining; Blue: Hoechst (Nucleus marker); Green: ATP7A; Red: Giantin. See Figure S7B for images of other conditions. Representative images are shown. Bars = 10 µm. n=3 independent experiments. A zoomed inset is displayed on the outmost right panel. Number of cells analyzed ≥ 27 for each condition. A parametric ordinary one-way ANOVA test was carried out with a Dunnett’s Multiple Comparison Test **** p<0,00001.

Having validated that Cu(II)(DapHH-αR5W4^NBD^) and Cu(II)(HDapH-αR5W4^NBD^) i) can penetrate into PC12 cells and ii) localize inside vesicular structures, and iii) may gradually release Cu within endosomes, we then indirectly probed the bioavailability of the imported Cu by monitoring the delocalization of the Golgi-associated Cu transporter, ATP7A. This protein is known to delocalize from the perinuclear Golgi area to vesicles in proximity to the plasma membrane when the cell needs to expel Cu in excess^48^. Treatment with Cu(II)(HDapH-αR5W4^NBD^) induced a significant delocalization of ATP7A to vesicular structures close to the plasma membrane in PC12 cells after 1h incubation, thereby confirming the presence of cytosolic bioavailable Cu (Figure 2B and S7B). However, Cu(II)(DapHH-αR5W4^NBD^) only showed a tendency toward ATP7A delocalization, which is in line with the slower increase in fluorescent signal in these cells (Figure 2A). These observations suggest that although DapHH-αR5W4^NBD^ is internalized into the cell, it is slow to release bioavailable Cu. It is of note that the shuttles are not fixable and therefore do not contribute to the signal of ATP7A in green when cells were fixed with PFA, as confirmed by controls (Figure 2B and S7B).

Next, we investigated the ability of DapHH-αR5W4^NBD^ and HDapH-αR5W4^NBD^ to protect PC12 from Cu(II)-Aβ peptides induced toxicity in the presence of a physiologic concentration of AscH^-^ (100 µM)^49–51^. Cu(II)Aβ_1-16_ was toxic only in the presence of the reductant. This toxicity was reversed in the presence of DapHH-αR5W4^NBD^ or HDapH-αR5W4^NBD^, but not in the presence of AKH-αR5W4^NBD^ (Figure 3A), except after preincubation of AKH-αR5W4^NBD^ with Cu(II)Aβ_1-16_ for an hour. This is in line with the faster retrieval of Cu(II) from Aβ and arrest of ROS production by the DapHH-αR5W4^NBD^ and HDapH-αR5W4^NBD^ shuttles as demonstrated *in vitro* (Figure 1C,D). The ability of these shuttles to rescue PC12 cells from Cu-induced toxicity was also studied using three independent batches of the full-length Aβ_1-40_ peptide. In agreement with the known variability in Aβ-induced cell toxicity^52^, of the three batches of Aβ_1-40_ tested, only batch 1 and 2 had a significant toxicity on PC12 cells at 10 µM, whereas batch 3 showed only a tendency (Figure 3B, Figure S8). In the presence of Cu(II)Aβ_1-40_, upon addition of AscH^-^, there was a significant increase in toxicity for all three Aβ_1-40_ batches due to Cu(Aβ)-induced ROS production, as observed for Aβ_1-16_ (Figure 3A). Importantly, DapHH-αR5W4^NBD^ and HDapH-αR5W4^NBD^ fully rescued this toxicity. In conclusion, both DapHH-αR5W4^NBD^ and HDapH-αR5W4^NBD^, are capable of preventing Cu(II)Aβ_1-16_ and Cu(II)Aβ_1-40_ induced toxicity in PC12 cells in the presence of AscH^-^.

**Figure 3:**
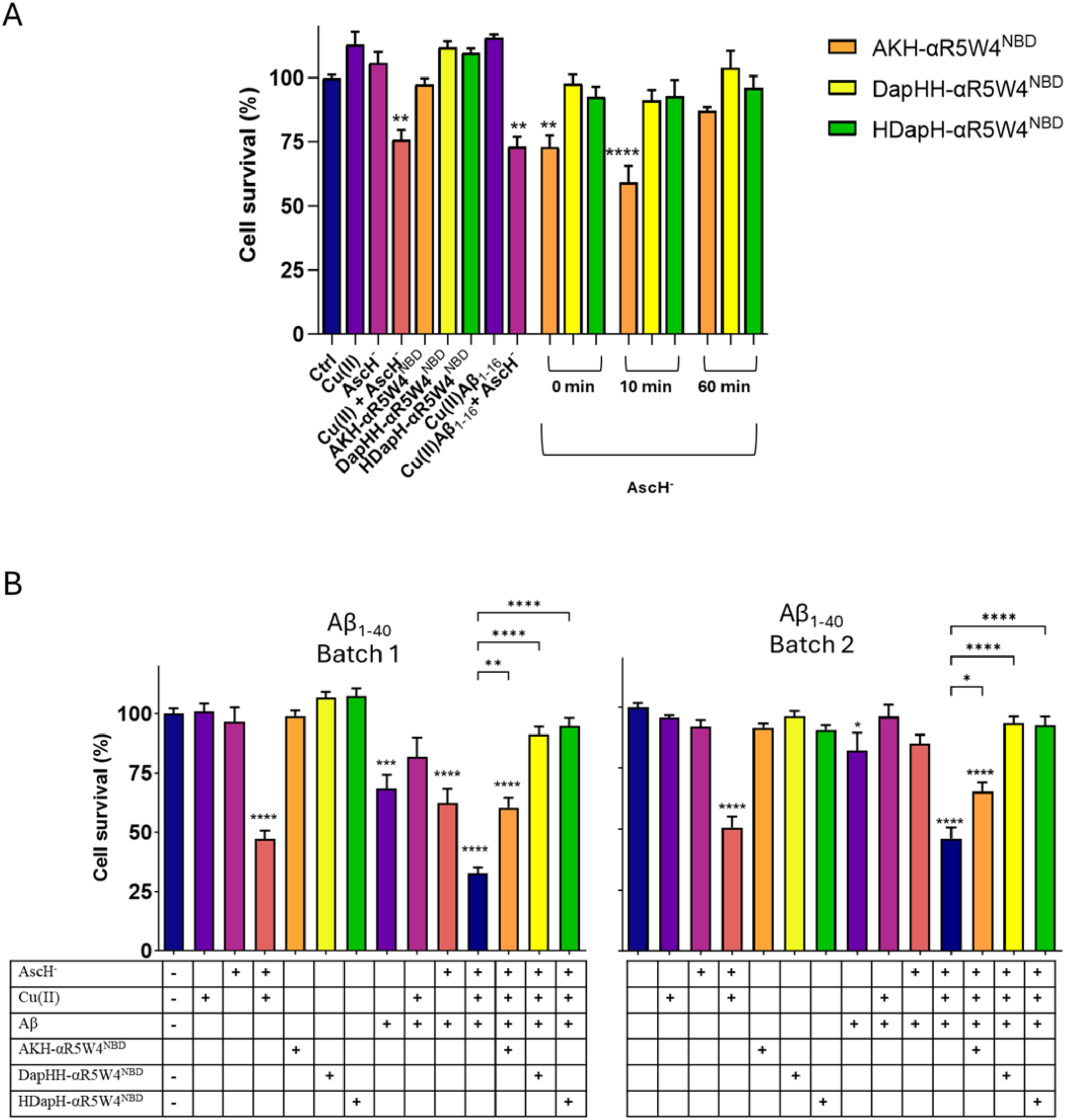
DapHH-αR5W4^NBD^ and HDapH-αR5W4^NBD^ prevent Cu-induced ROS and associated toxicity in PC12 cells. **(A**) 5 µM AKH-αR5W4^NBD^, DapHH-αR5W4^NBD^ or HDapH-αR5W4^NBD^ were incubated with 10 µM Cu(II)Aβ_1-16_ 0.5:1 in 10% DMEM in a test tube for 0, 10 or 60 min before addition of 500 µM AscH^-^ and immediate administration on the PC12 cells. Experiments were done in triplicates, n=3. Passed a normality D’Agostino & Pearson test. A parametric ordinary one-way ANOVA test was carried out with a Tukey Post Test * p<0,01, ** p<0,001, **** p<0,00001. (**B**) PC12 cells were incubated with 5 µM AKH-αR5W4^NBD^, DapHH-αR5W4^NBD^ or HDapH-αR5W4^NBD^ with 10 µM Cu(II)(Aβ_1-40_) 0.5:1 and 500 µM AscH^-^ for 24h. Experiments were done in triplicates, n=3. A parametric ordinary one-way ANOVA test was carried out with a Tukey’s multiple comparison Test * p<0,01, ** p<0,001, **** p<0,00001. Experiments were carried out in 10% DMEM. “+” signifies presence of a particular molecule.

### Organotypic hippocampal slice culture (OHSC) model of AD

OHSC is a relevant model to study neurodegenerative diseases, and has been used to investigate Aβ toxicity^53,54,55,56^, oxidative stress remediation^57,58^ and glial activation^59,60^ linked to Alzheimer’s disease. Its value lies in its similarity to *in vivo* animal models, particularly the preservation of neuronal connections. Therefore, after establishing the shuttles’ ability to prevent ROS production, to penetrate cellular membrane and deliver bioavailable Cu in a monoculture system, we next sought to determine whether these complex functions, translated to an overall tissue protection in an OHSC model of extracellular Cu-driven toxicity in the context of AD.

Brain slices with a visible dendate gyrus (DG), CA3, CA2, and CA1 regions were obtained from CD1 mice between P6-P10 and placed in culture for 2 weeks before treatment. Hippocampal slices were incubated with Cu(II)Aβ_1-40_ or Cu(II)Aβ_1-42_ in the presence or absence of DapHH-αR5W4^NBD^or HDapH-αR5W4^NBD^ for 48h without AscH^-^ supplementation because the brain intrinsically possesses between 0.1 to 1 mM AscH^-49,50^, in contrast to cultured 2D cells^61^.

Cell density was analyzed by counting the number of Hoechst-labeled nuclei, which were similar for each condition validating slice quality and indicating low variability (Figure 4A-B). Cell toxicity on organotypic hippocampal slices was determined by treating slices for 1h with propidium iodide (PI) before Hoechst staining. PI-labelled cells represent cells with a compromised plasma membrane thus permitting PI to intercalate in to cell DNA. Of note, PI staining does not directly detect cell death, but rather permeabilization of the plasma membrane, one the first stages toward cell death upon ROS-induced membrane alteration.

**Figure 4:**
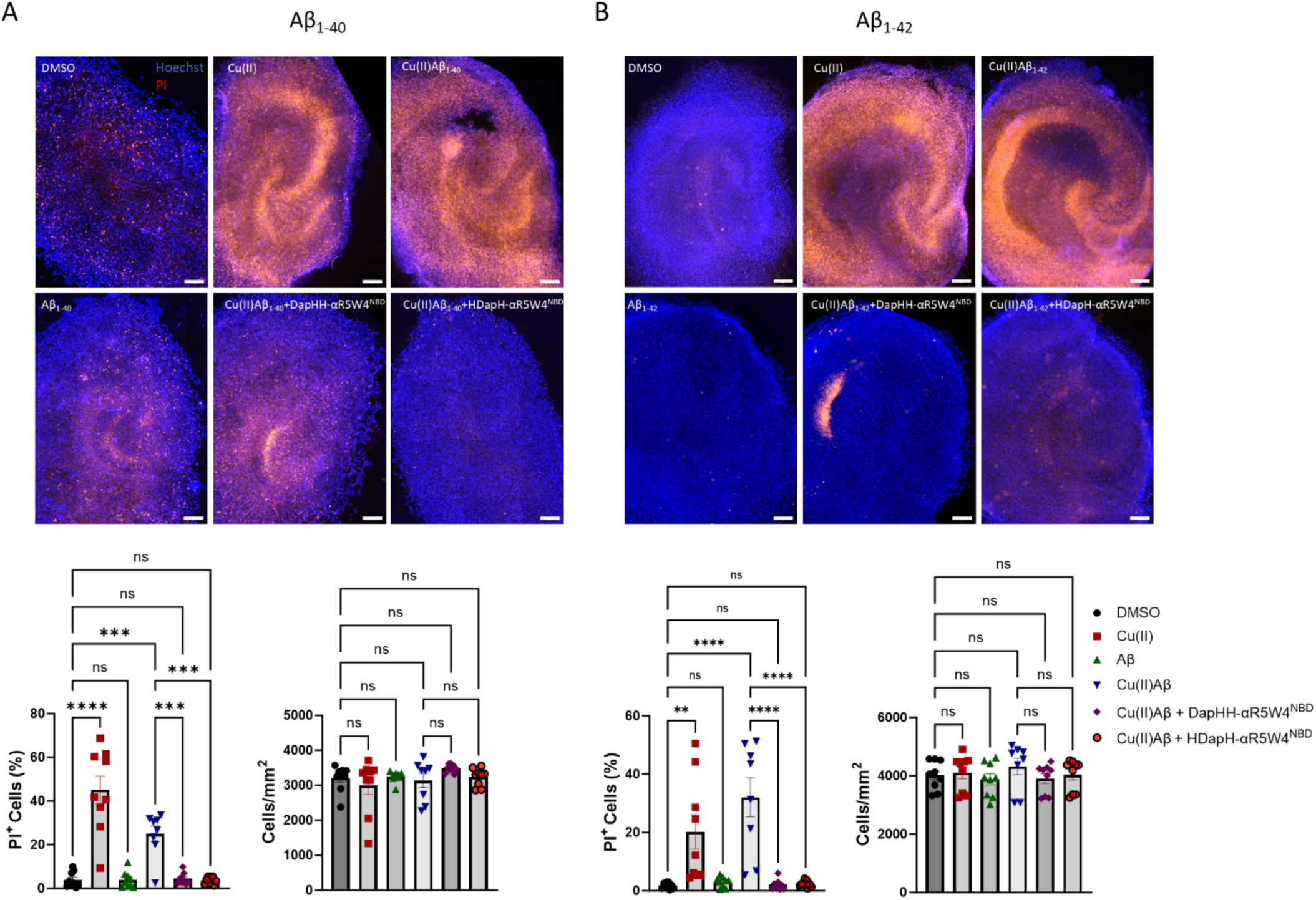
DapHH-αR5W4^NBD^ or HDapH-αR5W4^NBD^ rescue Cu(II) induced cell toxicity in OHSC. **(A)** Effect of Cu(II)(Aβ_1-40_) or **(B)** Cu(II)(Aβ_1-42_) on OHSC toxicity. Representative images of the effect of Cu(II) peptide shuttles on Cu(II)(Aβ_1-40/42_)-induced toxicity on OHSC with analysis of the preventive effect of DapHH-αR5W4^NBD^ or HDapH-αR5W4^NBD^ and analysis of the cell density in OHSCs culture after 48h treatment. Conditions: 10 µM Cu(II), DapHH-αR5W4^NBD^ or HDapH-αR5W4^NBD^=, 20 µM Aβ_40_ or Aβ_42_. Hoechst is blue and PI red. Scale: 200µm. A non-parametric test was carried out with a Kruskal-Wallis Multiple Comparison Test for Cells/mm^2^ for both Aβ_40_ and Aβ_42_. A parametric ordinary one-way ANOVA test was carried out with a Tukey’s Multiple Comparison Test for PI^+^ cells for both Aβ_1-40_ and Aβ_1-42_,** p<0,001, *** p<0,0001, **** p<0,00001. n=3 independent experiments with at least 3 slices per condition.

Treatment of OHSCs with 20 µM of monomeric Aβ_1-40_ or Aβ_1-42_ for 48h did not lead to detectable toxicity (Figure 4). However, in Cu(II) treated cells, there was a significant increase in PI-stained nuclei, suggesting that 10 µM of extracellular Cu(II) is toxic to organotypic hippocampal slices. Accordingly, 10 µM Cu(II) activated microglia in primary cell culture, which may also trigger neuronal damage^62^. Potential mechanisms for this Cu-induced toxicity include neuronal Cu overloading, Cu(Aβ)-induced ROS production in line with the endogenous presence of AscH^-^ and microglial activation by “loosely”-bound extracellular Cu. Treatment with Cu(II)(Aβ_1-40_) or Cu(II)(Aβ_1-42_) also triggered an increase in slice toxicity, which could be attributed to similar mechanisms as those described for Cu(II) (Figure 4, S9). Importantly, co-incubation with Cu(II)(Aβ_1-40/42_) and DapHH-αR5W4^NBD^ or HDapH-αR5W4^NBD^ completely rescued this toxicity. Together, these results are in line with our findings in PC12 cells and also reinforce the hypothesis that the Cu(II)(Aβ_1-40/42_)-induced toxicity arises from an extracellular effect. In other words, the toxicity observed in our assays is probably not the consequence of an increase in intracellular Cu levels, which is expected to be further increased by these Cu(II)-selective shuttles, but rather results from Cu(II)(Aβ_1-40/42_) ROS induction, which is rescued by the shuttles.

Proliferation and overactivation of microglia in the brain, concentrated around amyloid plaques, is a prominent feature of AD^63^. Given that Cu(II) could induce microglia activation both directly or through ROS production^62,64,65^, treated slices were stained using Iba1, a pan microglia marker^66^ and CD68, a marker of activated microglia^67^. Treatment of OHSC with Cu(II), Cu(II)Aβ_1-40_ or Cu(II)Aβ_1-42_ induced an increase in the percentage of microglial cells compared to DMSO-treated control slices (Figure 5). Since cell density remained unchanged after treatment with Cu(II)Aβ_1-40/42_ (Figure 4), this could signify that the induced proliferation of microglial cells occurs concomitantly with death of other cell types in Cu(II) treated OHSC slices. Importantly, co-incubation with DapHH-αR5W4^NBD^ or HDapH-αR5W4^NBD^ reduced microglial cell proliferation induced by Cu(II)(Aβ_1-40/42_) (Figure 5). Equally, there was an increase in microglial activation in Cu(II), Cu(II)Aβ_1-40_ or Cu(II)Aβ_1-42_ treated slices, as seen by an increase in CD68 labelling in microglial cells (Iba1 and CD68 positive cells), which was efficiently prevented by co-incubating Cu(II)(Aβ_1-40/42_) treated slices with the Cu(II)-selective shuttles. In conclusion, DapHH-αR5W4^NBD^ and HDapH-αR5W4^NBD^ display protective activity towards brain tissue against Cu(II) and Cu(II)(Aβ_1-40/42_)-induced insult.

**Figure 5:**
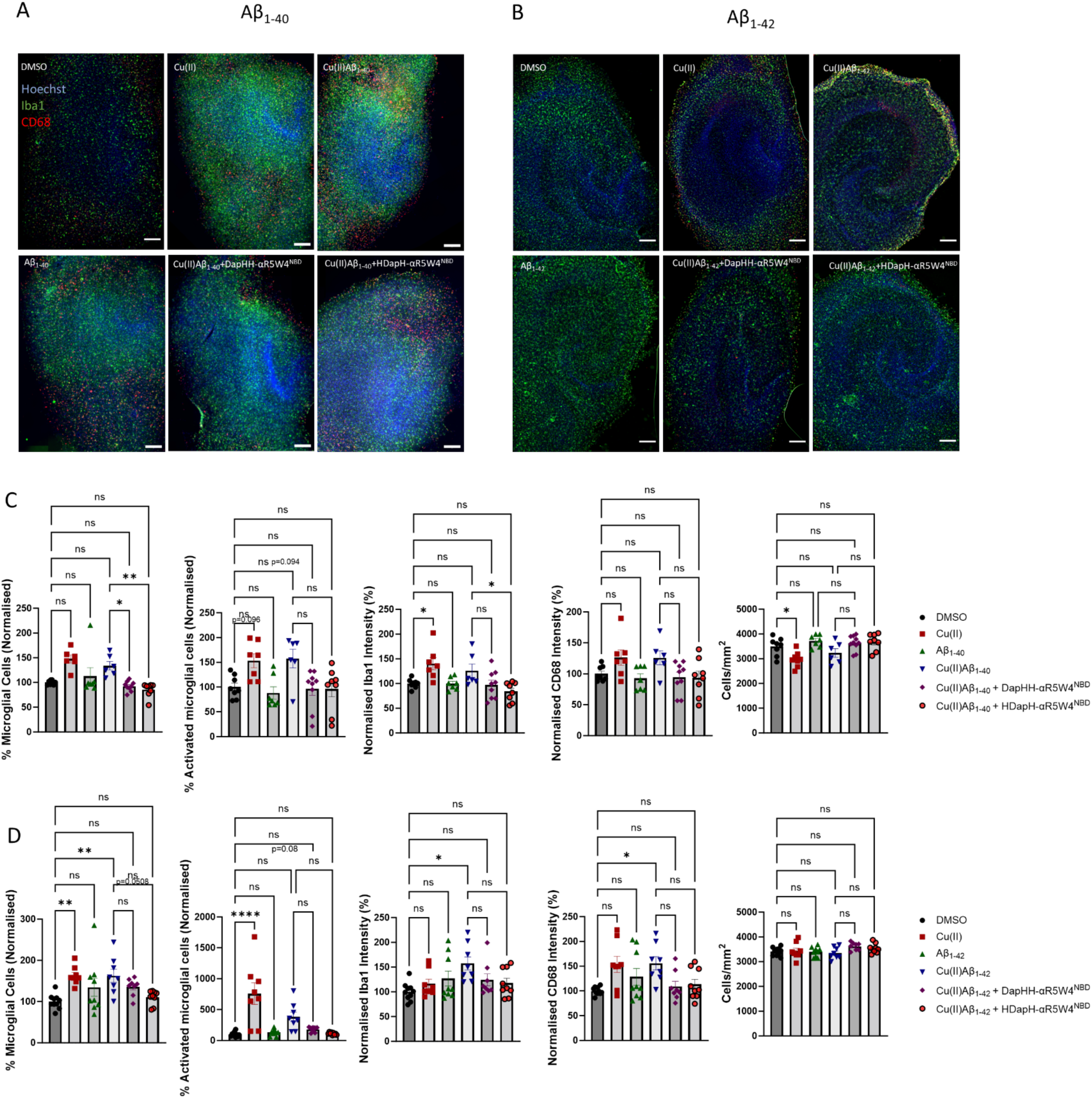
Effect of DapHH-αR5W4^NBD^ or HDapH-αR5W4^NBD^ on Cu(II)-induced microglia proliferation and activation. (A-B) Representative images of the effect of Cu(II) peptide shuttles on Cu-induced Cu(II)Aβ_40/42_ microglia activation on OHSC with **(C-D)** analysis of the preventive effect of DapHH-αR5W4^NBD^ or HDapH-αR5W4^NBD^ on Cu(II)(Aβ_1-40/42_) induced microglia proliferation, activation, and analysis of the cell density in OHSCs culture after 48h treatment. Hoechst is blue, Iba1 green and CD68 red. Scale: 200µm. Conditions: 10 µM Cu(II)=DapHH-αR5W4^NBD^ or HDapH-αR5W4^NBD^, 20 µM Aβ_1-40/42_. A non-parametric test was carried out with a Kruskal-Wallis Multiple Comparison Test for activated microglial cells and % microglial cells for Aβ_1-40_ and normalized Iba1 intensity for Aβ_1-42_. A parametric ordinary one-way ANOVA test was carried out with a Tukey’s Multiple Comparison Test for the rest, * p<0,01,** p<0,001, *** p<0,0001, **** p<0,00001. n=3 independent experiments with at least 3 slices per condition.

## Discussion

Cu has been studied for its role in many pathologies such as Wilson and Menkes diseases as well as several neurodegenerative disorders including AD^68–79^. Although the origins of Cu-related dysfunction have been elucidated in some cases (*e. g.* Wilson’s disease and Menkes disease), many open questions remain, particularly concerning the mechanisms leading to these Cu dyshomeostasis and how the loss of Cu homeostasis affects different organs, especially the brain^80^. In this context, tools to study Cu-metabolism from cell culture to *in vivo* models are crucial to better delineate intrinsic mechanisms. To date only a few studies have employed Cu(II)-selective fluorescent sensing probes capable of entering cells^81,82^. Although the molecules used in those studies cross cell membranes, they either display a very weak Cu(II)affinity, or lack specificity, in contrast to the shuttles developed in the present work. A major advantage of the shuttles developed here is their ability to be tracked in real-time within cells. This is due to the positioning of a fluorophore close to the Cu(II) coordination sphere, which permits up to 98% fluorescence quenching upon Cu(II) binding (Figure S2A). Hence, restoration of the shuttle fluorescence upon intracellular Cu release provides both temporal and subcellular resolutions of the Cu release process, making AKH-αR5W4^NBD^, DapHH-αR5W4^NBD^ and HDapH-αR5W4^NBD^ particularly well-suited for mechanistic studies to follow Cu trafficking in cells.

Besides this first application, the shuttles described here were successfully designed to exhibit fast Cu(II)-binding kinetics, enabling efficient Cu(II) retrieval rate from Aβ (Figure 1C, D). This property is essential, because it permits the retrieval of extracellular Cu from Aβ, before internalization into cells. The importance of this parameter is demonstrated in this study, as DapHH-αR5W4^NBD^ and HDapH-αR5W4^NBD^ immediately suppressed Cu(Aβ)-induced ROS production immediately and protect both cells and tissue against Cu(II)-Aβ induced toxicity, in contrast to AKH-αR5W4^NBD^ (Figures 3 and 4). Importantly, in our OHSC model, DapHH-αR5W4^NBD^ and HDapH-αR5W4^NBD^ were able to reverse the increased proliferation and activation of microglial cells after treatment with Cu(II) or Cu(II)(Aβ) (Figure 5), both believed to be associated with brain toxicity provoked by Cu-induced ROS production^67,83,84^. Notwithstanding, our findings only demonstrate protection against Cu-induced injury and cannot be directly extrapolated to Aβ-driven neurodegeneration, as short time incubation with monomerized Aβ peptides alone did not induce toxicity in our OHSC model. Future studies in transgenic OHSC models that exhibit intrinsic Aβ toxicity will be essential.

Other important characteristics for good Cu shuttle are their intracellular kinetics, Cu release mechanism and intracellular location of Cu release. For instance, we showed here that nanomolar concentrations of Cu(II)GTSM (Figure S6B), was toxic to PC12 cells, and the toxicity of Clioquinol have been previously reported^85^. It has equally been shown that Cu ionophores such as Elesclomol, induced the so-called cuproptosis at low nanomolar concentrations that was attributed to the Cu-induced aggregation of lipoylated mitochondrial proteins involve in Krebs cycle^86^. These features may partially explain the failure of Cu ionophores, such as Clioquinol and PBT2 in clinical trials against AD, in addition to their lack of Cu(II) selectivity^3,87^. Another factor to be considered is the delivery site of these ionophores, which have tendencies to accumulate Cu in the mitochondria. The accumulation of Cu in the mitochondria might compete with other transition metal ions (mainly Fe) and provoke mitochondrial dysfunction as well as cuproptosis^88^. Hence, we expect that a slow and gradual release of Cu(II) from endolysosomal vesicles might be more beneficial and less toxic than fast cytosolic accumulation of Cu which might overwhelm the cellular metal ion buffering system^89^. Subsequently, the imported Cu can exit endosomes via CTR1/CTR2 or DMT1 transporters^90,91^, making the Cu bioavailable as seen by ATP7A delocalization (Figure 2B).

In line with this, we further hypothesized that the Cu-release from the Cu-shuttle described here occurs via thiol (ex: Cystine or GSH) reduction in endosomes. Although reduction by GSH was effective *in vitro* (Figure 1C), *in cellulo* data indicated that reduction alone is probably not the main mechanism at play. Indeed, we observed that AKH-αR5W4^NBD^ and HDapH-αR5W4^NBD^ release Cu quicker into cells than DapHH-αR5W4^NBD^ (Figure 2B, S7B), while the latter is the fastest to be reduced by GSH *in vitro*. Hence, an alternative but not mutual exclusive hypothesis may link Cu release by the shuttles to a progressive acidification along the endolysosomal pathway, the major cellular entry routes used by αR5W4 CPP^39^, or a combination of acification and reduction. Indeed, the XxxHisHis motif binds Cu(II) down to pH 5,^44,45,92,93^ with a conditional affinity of 10^7^ M^-1^ corresponding to about 95% of bound Cu(II) at 10 µM^93^. In contrast, the HisXxxHis motif releases Cu(II) at higher pH, analogously to XxxZzzHis that has a lower conditional affinity (10^5^ M^-1^) leading to only ∼50% of bound Cu(II)^94^. Hence, this suggests that Cu(II) release from AKH-αR5W4^NBD^ and HDapH-αR5W4^NBD^ may occur earlier in the endolysosomal compartment (pH ∼ 5.5-6.5), whereas Cu(II) release by DapHH-αR5W4^NBD^ could occur only at the lower pH of lysosomes (pH ∼ 4.5-5.5).

## Conclusion

In conclusion, the Cu(II) peptide shuttles DapHH-αR5W4^NBD^ and HDapH-αR5W4^NBD^ bind Cu(II) with high affinity and selectivity and with sufficiently fast kinetic to supress Cu(Aβ)-induced ROS production. These peptide shuttles efficiently retrieve extracellular Cu(II) from Aβ and import Cu into neuronal cells, displaying distinct kinetics profiles of Cu(II) extraction and release (AKH-αR5W4^NBD^: Slow binding and fast release; DapHH-αR5W4^NBD^: intermittent binding and slow release; HDapH-αR5W4^NBD^: Fast binding and intermittent release). The Cu released within cells was further shown to be bioavailable and nontoxic. Importantly, these Cu(II) peptide shuttles also prevented Cu-and Cu(Aβ)-induced neurotoxicity, microglial proliferation and activation in OHSC. Therefore, they pose as promising drug candidates for future *in vivo* studies in AD animal models and could lead to therapeutic strategies aimed at correcting Cu-dyshomeostasis in patients with deregulated Cu levels.

## Acknowledgements

We acknowledge the Plateforme Imagerie In Vitro of ITI Neurostra at CNRS UAR-3256. Special thanks to Fréderic Doussau for his help with OHSC.

The authors declare that they have no conflicts of interest with the contents of this article.

## Data availability

Data availability can be requested to the corresponding authors.

