## Supplemental Figures for "Dual-action peptide shuttles rescue Cu-amyloid-β-induced neurotoxicity and relocate Cu intracellularly"

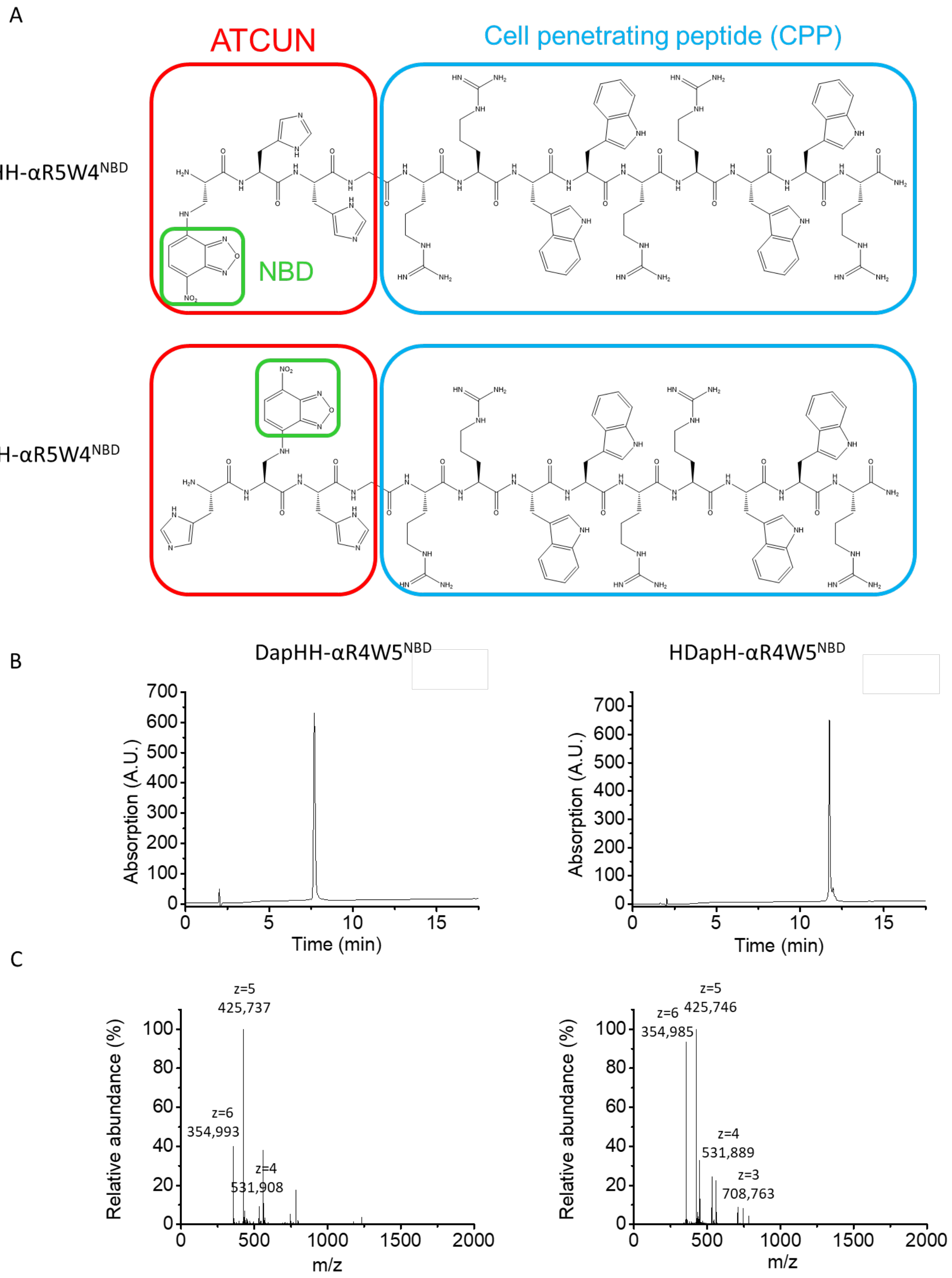

A
ATCUN
Cell penetrating peptide (CPP)
DapHH-αR5W4NBD
NBD
HDapH-αR5W4NBD
B
DapHH-αR4W5NBD
HDapH-αR4W5NBD
Absorption (A.U.)
Time (min)
C
Relative abundance (%)
m/z
z=5
425,737
z=6
354,993
z=4
531,908
z=6
354,985
z=5
425,746
z=4
531,889
z=3
708,763

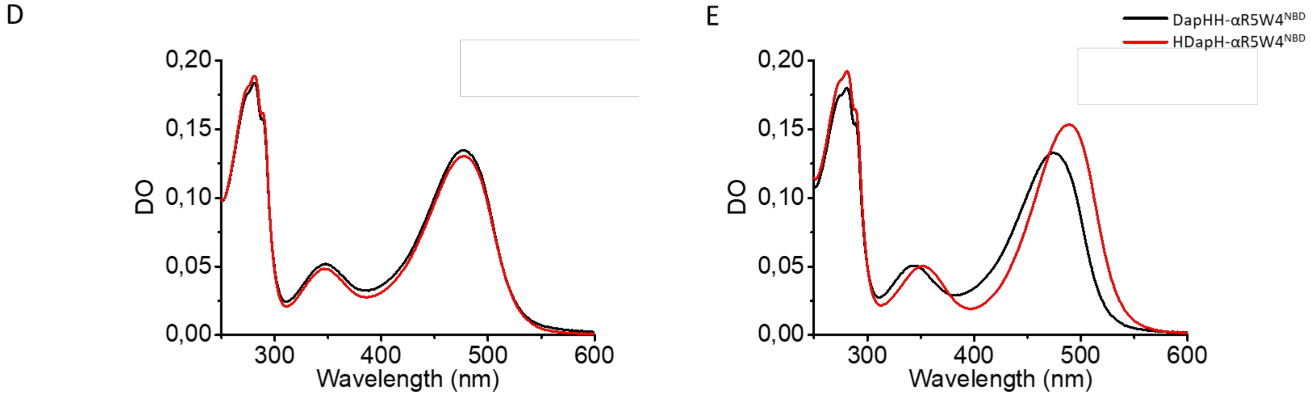


Figure S1: (A) Amino acid sequence of DapHH-αR5W4NBD and HDapH-αR5W4NBD. (B) HPLC chromatogram of purified DapHH-αR4W5$^{NBD}$ and HDapH-αR4W5$^{NBD}$. The separation was performed using a linear gradient of buffer A (TFA 0.1%) and buffer B (TFA 0.1%, ACN 90%) ranging from 5% buffer B to 100% buffer B in 30 min, flow 1 ml/min, and UV detection at 214 nm. (C) MS of major peak in LC spectra of purified DapHH-αR4W5$^{NBD}$ and HDapH-αR4W5$^{NBD}$. The separation was performed using a linear gradient of buffer A (Formic acid 0.1%) and buffer B (Formic acid 0.1%, ACN 90%) ranging from 5% buffer B to 100% buffer B in 15 min, flow 1 ml/min, and m/z detection from 100 to 2000 Da. (D) UV-Vis spectra of DapHH-αR5W4$^{NBD}$ and HDapH-αR5W4$^{NBD}$ in 100 mM HEPES or (E) Cu(II)DapHH-αR5W4$^{NBD}$ and Cu(II)HDapH-αR5W4$^{NBD}$. Conditions: Cu(II)DapHH-αR5W4$^{NBD}$=Cu(II)HDapH-αR5W4$^{NBD}$= 5 µM, HEPES 100 mM pH 7,4, 25°C. Representative traces of n=2 independent experiments are shown.

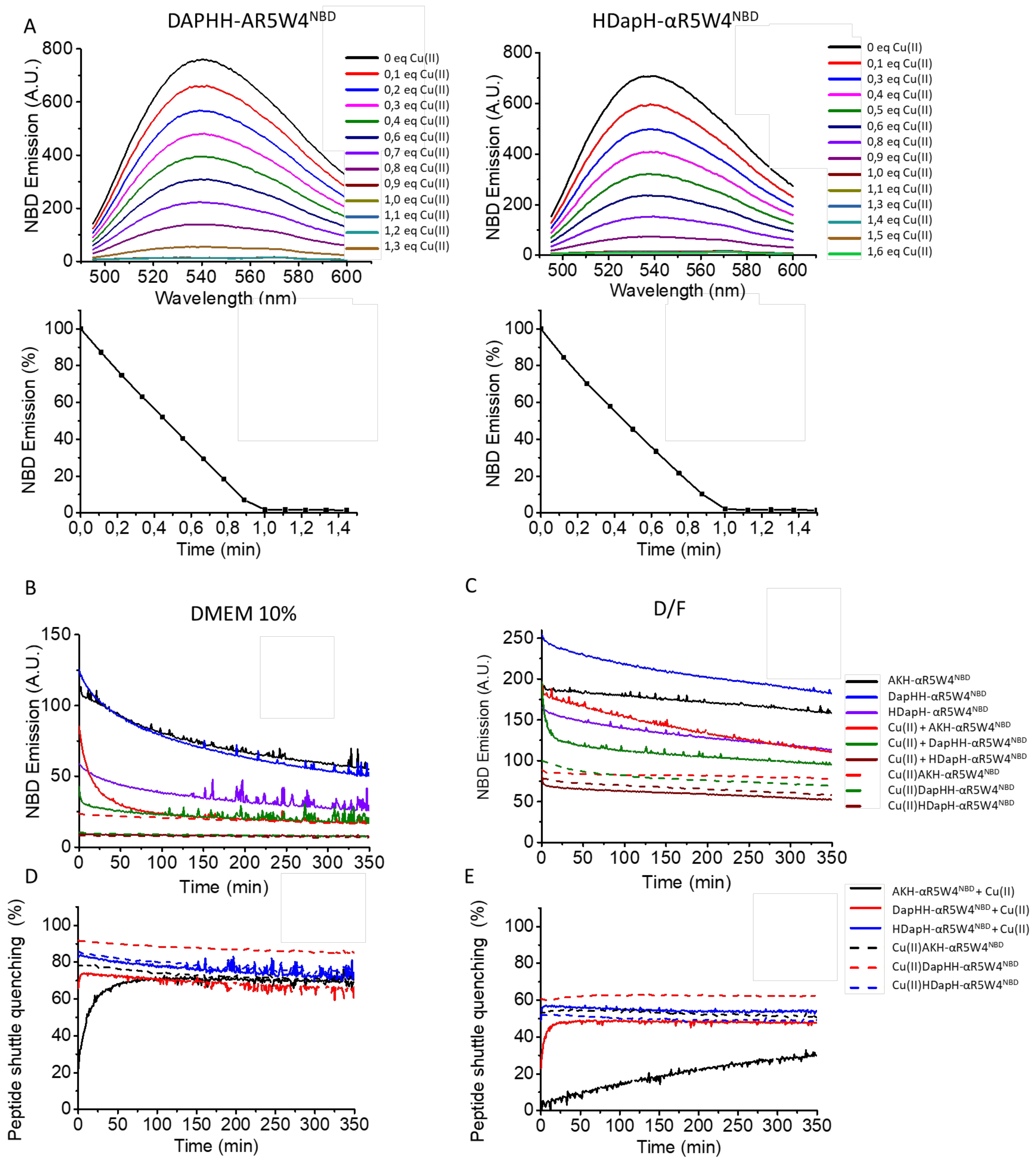


Figure S2: (A) Fluorescence spectra of DapHH-αR5W4$^{NBD}$ and HDapH-αR5W4$^{NBD}$ along with their respective linear Cu(II) dependent quenching. Spectra taken after additions of 0.1 equivalent Cu(II) and fluorescence intensity was monitored at 545 nm. Conditions: 4 µM peptides shuttles, addition of Cu(II) ; 0.1 equivalent = 0.4 µM ; HEPES Buffer

100 mM pH 7.4. Excitation wavelength: 477 nm; Emission spectra: 490-600 nm. Representative traces of n=2 independent experiments are shown. (B) Withdrawal of Cu(II) by peptide shuttles in cell culture media. NBD emission of AKH-αR5W4$^{NBD}$, DapHH-αR5W4$^{NBD}$ or HDapH-αR5W4$^{NBD}$ in DMEM 10% or (C) D/F, in the presence or absence of Cu(II), monitored at 545 nm. Dashed lines represent the emission of peptide shuttles precomplexed to Cu(II), thus the maximum theoretical quench. (D) Normalised percentage of AKH-αR5W4$^{NBD}$, DapHH-αR5W4$^{NBD}$ or HDapH-αR5W4$^{NBD}$ in DMEM 10% (E) or D/F, in the presence or absence of Cu(II) at 545 nm. Dashed lines represent the emission of peptide shuttles precomplexed to Cu(II), thus the maximal theoretical quench (See Formula S1 below). Conditions: Cu(II)=AKH-αR5W4$^{NBD}$=DapHH-αR5W4$^{NBD}$=HDapH-αR5W4$^{NBD}$= 5 µM, DMEM 10%, D/F 100%, 25°C. Representative traces of n=3 independent experiments are shown.

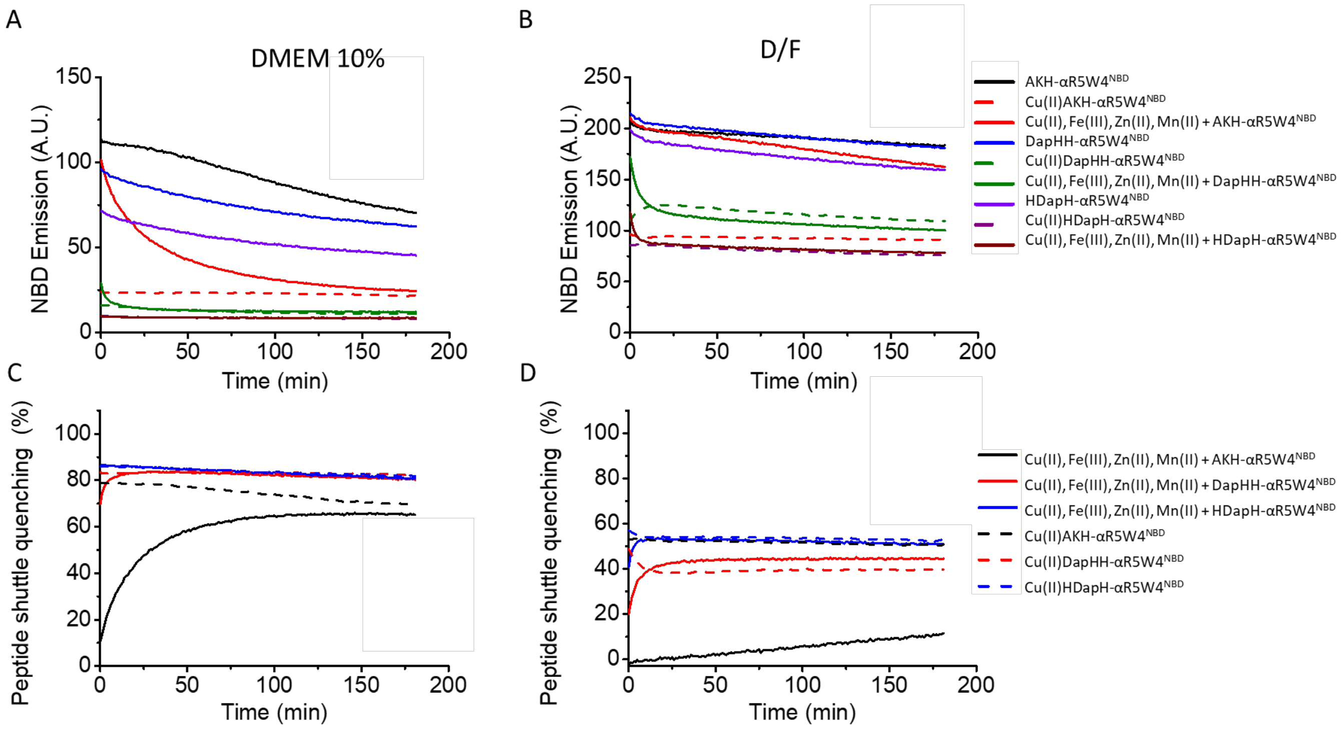


Figure S3: Selective withdrawal of Cu(II) by peptide shuttles in cell culture media containing Cu(II), Fe (III), Zn(II) and Mn(II). (A) NBD emission of AKH-αR5W4$^{NBD}$, DapHH-αR5W4$^{NBD}$ or HDapH-αR5W4$^{NBD}$ in DMEM 10% (B) or D/F, in the presence or absence of Cu(II) at 545 nm. Dashed lines represent the emission of peptide shuttles precomplexed to Cu(II), thus the maximum theoretical quench. (C) Normalised percentage of AKH-αR5W4$^{NBD}$, DapHH-αR5W4$^{NBD}$ or HDapH-αR5W4$^{NBD}$ in DMEM 10% (D) or D/F, in the presence or absence of Cu(II) at 545 nm. Dashed lines represent the emission of peptide shuttles precomplexed to Cu(II), thus the maximal theoretical quench (See Formula S1 below). Conditions: Cu(II)=AKH-αR5W4$^{NBD}$=DapHH-αR5W4$^{NBD}$=HDapH-αR5W4$^{NBD}$= 5 µM, DMEM 10%, D/F 100%, 25°C. Representative traces of n=3 independent experiments are shown.

$$Cu(II)\ (A) + XZH\text{-}\alpha R5W4^{NBD}\ (B) \rightleftharpoons Cu(II)XZH\text{-}\alpha R5W4^{NBD}\ (C)$$

$$\left\{1 - \frac{(C)_t}{(B)_t}\right\} \times 100 = \text{maximal theoretical quench (\%)}$$

$$\left\{1 - \frac{(A+B)_t}{(B)_t}\right\} \times 100 = \text{\% Shuttle quenching at timepoint (t)}$$

Formula S1: Calculation of Peptide Shuttle quenching (%) for Figure S2D,E and Figure S3C,D. The maximal theoretical quench is derived by dividing the fluorescence emission of Cu(II)XZH-αR5W4$^{NBD}$ by the emission of XZH-αR5W4$^{NBD}$ peptide shuttles (Dashed lines). % Shuttle quenching at timepoint (t) is derived by dividing the fluorescence emission of Cu(II) + XZH-αR5W4$^{NBD}$ by the emission of XZH-αR5W4$^{NBD}$ peptide shuttles (Bold lines).

| | Cu(II) | | Cu(II) + Fe (III) + Zn(II) + Mn(II) | |
|---|---|---|---|---|
| | DMEM 10% (min) | D/F (min) | DMEM 10% (min) | D/F (min) |
| AKH-αR5W4$^{NBD}$ | 22.0 ± 2.4 | N/A | 40.3 ± 5.7 | N/A |
| DapHH-αR5W4$^{NBD}$ | <1 min | 4.3 ± 0.6 | <1 min | 5.5 ± 0.6 |
| HDapH-αR5W4$^{NBD}$ | <1 min | <1 min | <1 min | <1 min |

Table S1: Average half-time of Cu(II) withdrawal from DMEM 10% or D/F media, in the presence or absence of other d-block metals, by AKH-αR5W4$^{NBD}$, DapHH-αR5W4$^{NBD}$ or HDapH-αR5W4$^{NBD}$, n=3.

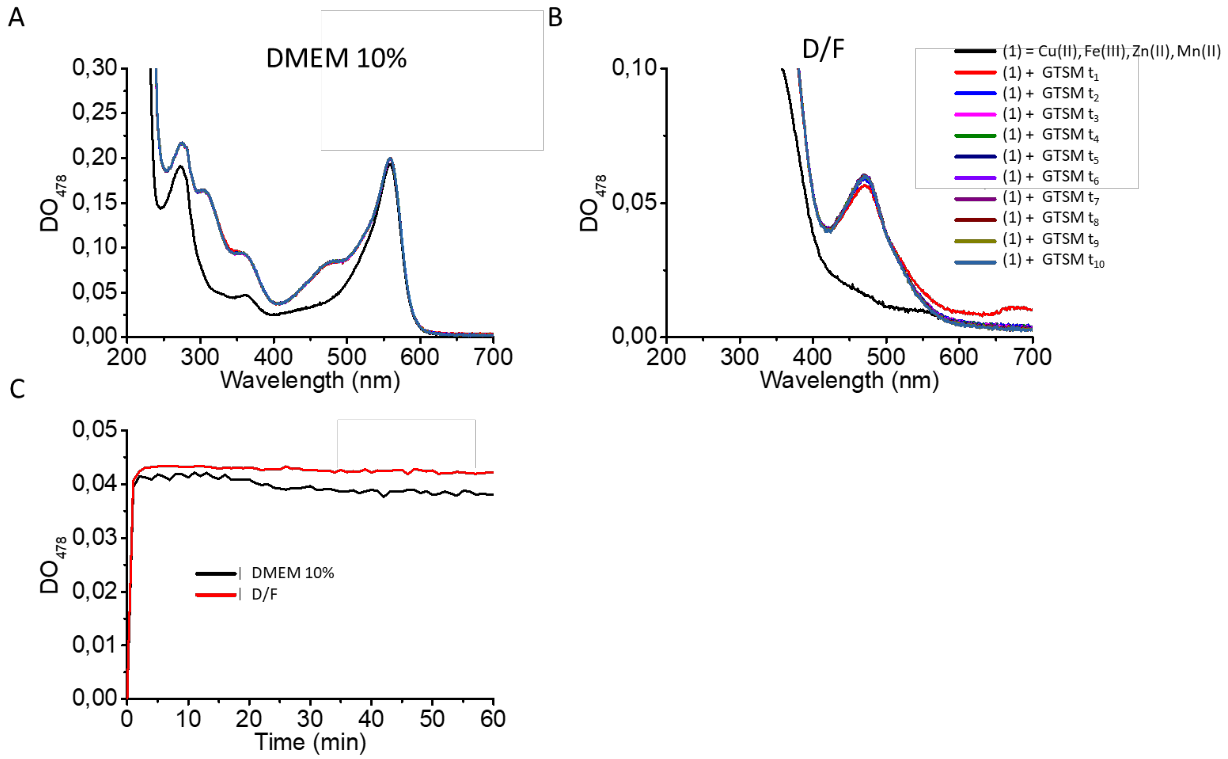


Figure S4: Selective withdrawal of Cu(II) by GTSM in cell culture media containing Cu(II), Fe(III), Zn(II) and Mn(II). d-d band absorption of Cu(II)GTSM in (A) DMEM 10% or (B) D/F media with an absorption maximum at 477 nm. (C) Net increase in absorption at 477 nm by Cu(II)GTSM complex in DMEM 10% vs D/F. Conditions: Cu(II)=GTSM= 5µM, DMEM 10%, D/F 100%, 25°C.

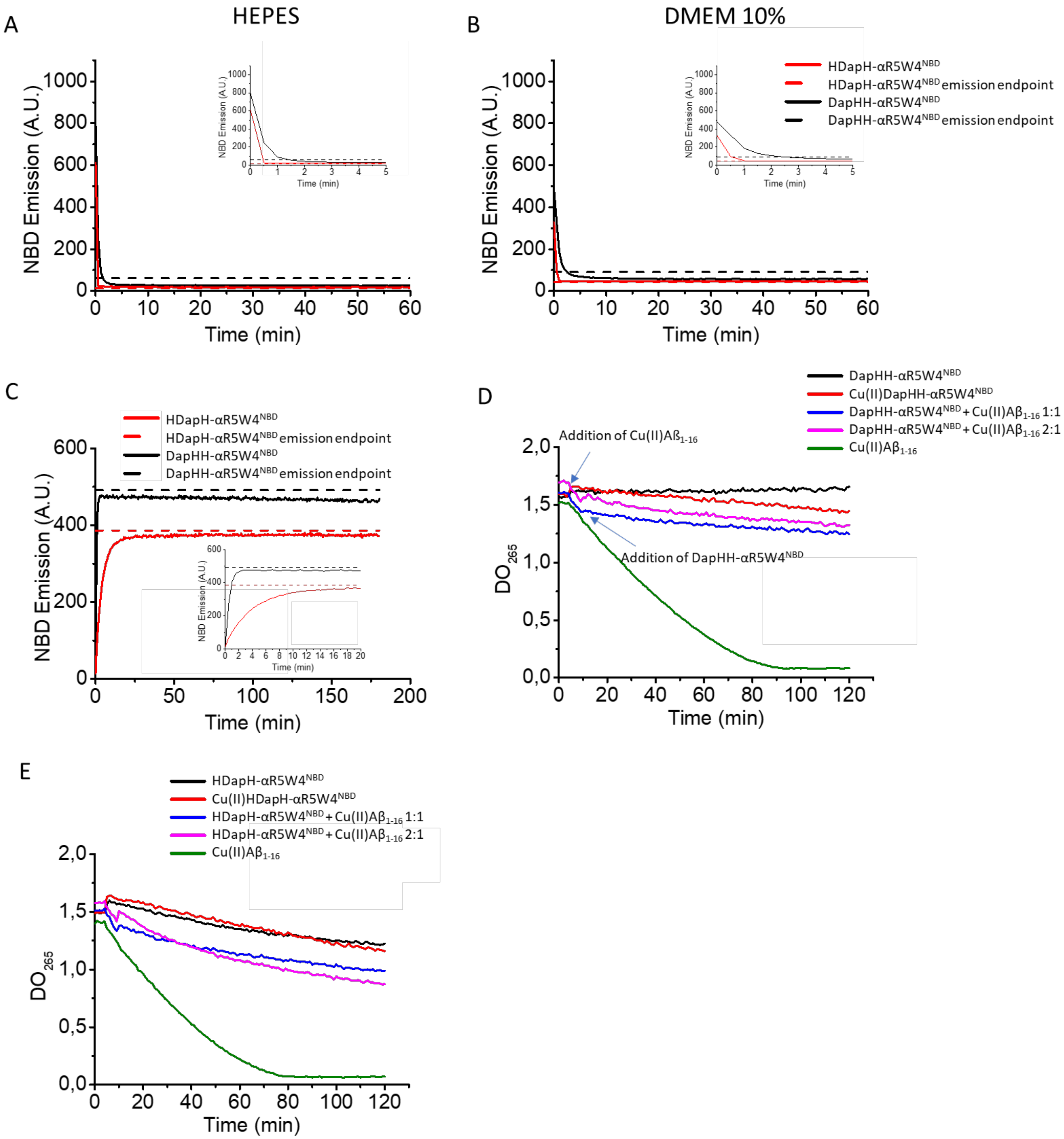


Figure S5: (A) Retrieval of Cu(II) from $A\beta_{1-16}$ by DapHH-αR5W4$^{NBD}$ (black line) and HDapH-αR5W4$^{NBD}$(red line), in HEPES or (B) DMEM 10% media. Dashed lines indicate the expected emission endpoint of DapHH-αR5W4$^{NBD}$ (black) and HDapH-αR5W4$^{NBD}$ (red), respectively, in the presence of Cu(II) in a 1:1 complex. Conditions: DapHH-αR5W4$^{NBD}$=HDapH-αR5W4$^{NBD}$= 5 µM; $A\beta_{1-16}$ =10 µM; Cu(II)=5 µM, HEPES 100 mM pH 7.4, 25°C. Representative traces of n=3 independent experiments are shown. (C) Reduction of Cu(II) bound to DapHH-αR5W4$^{NBD}$ and HDapH-αR5W4$^{NBD}$ by glutathione (GSH) monitored by the increase in NBD fluorescence at 545 nm over time. Black and red dashes, marks the expected emission endpoint (emission in the absence of Cu(II)) of DapHH-αR5W4$^{NBD}$ and HDapH-αR5W4$^{NBD}$, respectively. Conditions: Cu(II)DapHH-αR5W4$^{NBD}$=Cu(II)HDapH-αR5W4$^{NBD}$= 5 µM, 5 mM GSH,

HEPES 100 mM pH 7.4, 37°C; n=3 independent experiments. (D) Retrieval of Cu(II) from Aß$_{1-16}$ by DapHH-αR5W4$^{NBD}$ or (E) HDapH-αR5W4$^{NBD}$ halts ROS production. Inhibition of AscH$^-$ consumption by DapHH-αR5W4$^{NBD}$ or HDapH-αR5W4$^{NBD}$ monitored by the absorbance of AscH$^-$ at 265 nm. Addition of AscH$^-$ was carried out at $t_0$ and Cu(II)Aß1$_{-16}$ at $t_5$ and addition of XYH-αR5W4$^{NBD}$ (DapHH-αR5W4$^{NBD}$ or HDapH-αR5W4$^{NBD}$) at $t_{10}$. Conditions: AscH$^-$= 100 µM, Cu(II)DapHH-αR5W4$^{NBD}$=Cu(II)HDapH-αR5W4$^{NBD}$= 5 µM, Aß$_{1-16}$= 10 µM, Cu(II)= 5 µM, HEPES 100 mM pH 7,4. n=3 independent experiments.

| | Aß$_{1-16}$ | | GSH |
|---|---|---|---|
| | HEPES (s) | DMEM 10% (s) | HEPES (s) |
| DapHH-αR5W4$^{NBD}$ | 29.9 ± 5.2 | 47.9 ± 9.2 | 35.1 ± 1.9 |
| HDapH-αR5W4$^{NBD}$ | 6.6 ± 0.4 | 14.5 ± 3.8 | 241.9 ± 4.3 |

Table S2: Kinetics of Cu(II) transfer from Aß$_{1-16}$ to DapHH-αR5W4$^{NBD}$ and HDapH-αR5W4$^{NBD}$ in HEPES or 10% DMEM. Conditions: DapHH-αR5W4$^{NBD}$=HDapH-αR5W4$^{NBD}$=Cu(II)= 5 µM, Aß$_{1-16}$=10 µM, 100 mM HEPES pH 7,4, 10% DMEM, 25°C; n=3. (b) Reduction kinetics of Cu(II) on DapHH-αR5W4$^{NBD}$ or HDapH-αR5W4$^{NBD}$ by 5 mM GSH. Conditions: Cu(II)DapHH-αR5W4$^{NBD}$=Cu(II)HDapH-αR5W4$^{NBD}$= 5 µM, 5 mM GSH, HEPES 100 mM pH 7.4; 37°C; n=3 independent experiments.

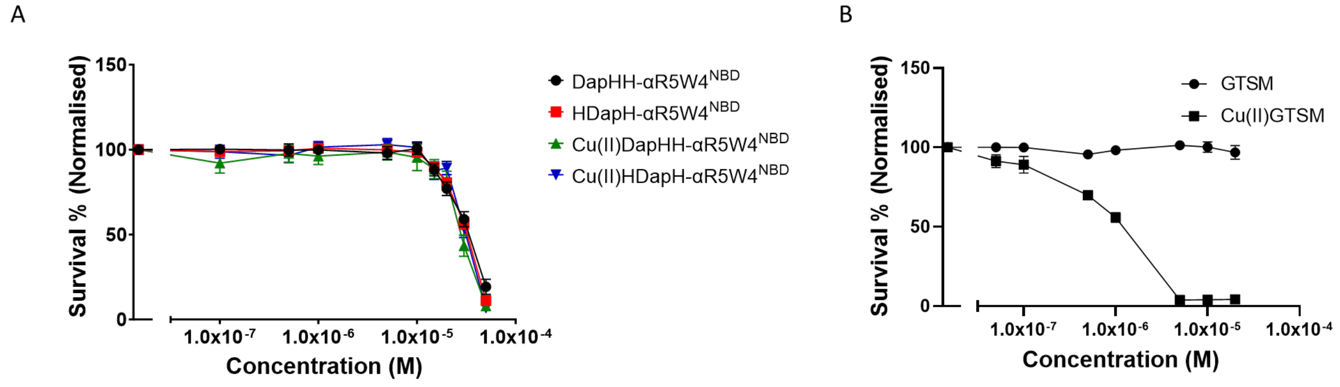


Figure S6: (A) Viability of PC12 cells after treatment with DapHH-αR5W4$^{NBD}$ or HDapH-αR5W4$^{NBD}$ alone or precomplexed with Cu(II). (B) Viability of PC12 cells after treatment with GTSM alone or precomplexed with Cu(II). Conditions: 5 x $10^4$ PC12 cells were incubated with the indicated concentration of peptides for 24h in 100% DMEM media.

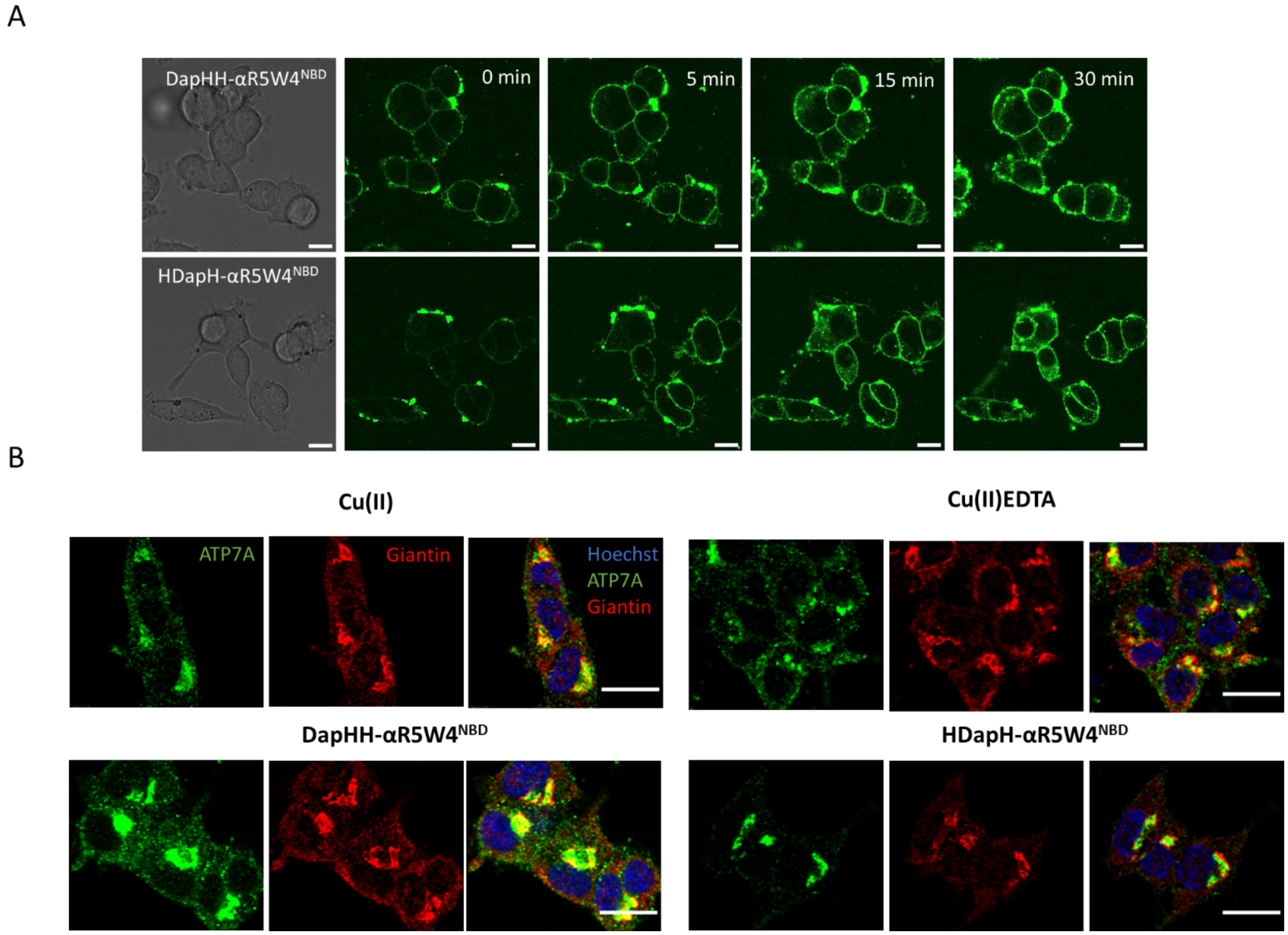


Figure S7: (A) Penetration of second-generation peptide shuttles into PC12 cells. Representative images at the indicated time obtained from live imaging performed on 5x10$^4$ PC12 cells incubated with 5 µM DapHH-αR5W4$^{NBD}$ or HDapH-αR5W4$^{NBD}$, at 37°C using a Leica TCS SP5 (II) confocal microscope, with an excitation wavelength at 477 nm for 30min. Time point zero is about 1 minute after incubation with the peptides. Bars = 10 µM. Similar observations were obtained with at least 3 independent cell passages. (B) Colocalization between ATP7A and Giantin staining. PC12 cells were incubated for 1h in DMEM media alone (control = Ctrl) or, 5 µM Cu(II), Cu(II)-EDTA, DapHH-αR5W4$^{NBD}$, Cu(II)DapHH-αR5W4$^{NBD}$, Cu(II)HDapH-αR5W4$^{NBD}$, HDapH-αR5W4$^{NBD}$ or 1 µM Cu(II)-GTSM, before fixation and processed for immunostaining; Blue: Hoechst (Nucleus marker); Green: ATP7A; Red: Giantin (Golgi marker). Representative images are shown. Bars = 10 µm. n=3 independent experiments.

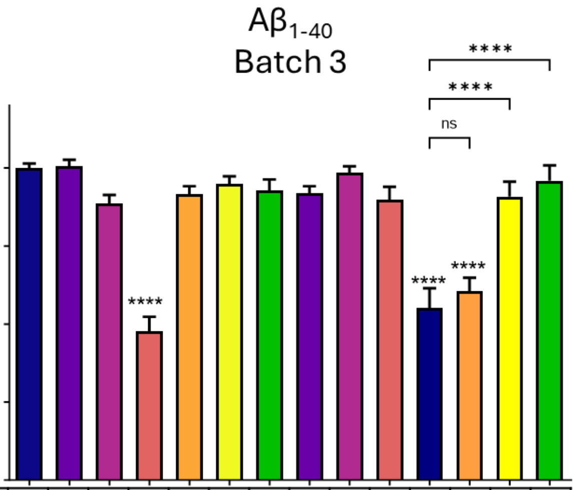


| | | | | | | | | | | | | | | |
|---|---|---|---|---|---|---|---|---|---|---|---|---|---|---|
| AscH- | | | + | + | | | | | | + | + | + | + | + |
| Cu(II) | | + | | + | | | | | + | | + | + | + | + |
| Aβ | | | | | | | | + | + | + | + | + | + | + |
| AKH-αR5W4$^{NBD}$ | | | | | + | | | | | | | + | | |
| DapHH-αR5W4$^{NBD}$ | | | | | | + | | | | | | | + | |
| HDapH-αR5W4$^{NBD}$ | | | | | | | + | | | | | | | + |

Figure S8: Transfer of Cu(II) from Aß$_{1-40}$ batch 3 to ATCUN motif prevents Cu-induced ROS production and toxicity to PC12 cells. PC12 cells were incubated with 5 µM AKH-αR5W4$^{NBD}$, DapHH-αR5W4$^{NBD}$ or HDapH-αR5W4$^{NBD}$ with 10 µM Cu(II)Aβ$_{1-40}$ 0.5:1 and 500 µM AscH$^-$ for 24h. Experiments were done in triplicates, n=3. A parametric ordinary one-way ANOVA test was carried out with a Tukey's multiple comparison Test * p<0,01, ** p<0,001, **** p<0,00001. Experiments were carried out in 10% DMEM. "+" signifies presence of a particular molecule.
